# PIWIL1 Promotes Gastric Cancer via a piRNA-Independent Mechanism

**DOI:** 10.1101/2020.05.03.075390

**Authors:** Shuo Shi, Zhen-Zhen Yang, Sanhong Liu, Fan Yang, Haifan Lin

**Affiliations:** Shanghai Institute for Advanced Immunochemical Studies, ShanghaiTech University, Shanghai, China 201210; Yale Stem Cell Center and Department of Cell Biology, Yale University School of Medicine, New Haven, CT 06519 USA

**Author notes:** Corresponding Author: Haifan Lin, email: < >.

**Keywords:** PIWI, gastric cancer, piRNA

## Abstract

Targeted cancer therapy aims to achieve specific elimination of cancerous but not normal cells. Recently, PIWI proteins, a subfamily of the PAZ-PIWI Domain (PPD) protein family, have emerged as promising candidates for targeted cancer therapy. PPD proteins are essential for small non-coding RNA pathways. The Argonaute subfamily partners with microRNA and small interfering RNA, whereas the PIWI subfamily partners with PIWI-interacting RNA (piRNA). Both PIWI proteins and piRNA are mostly expressed in the germline and best known for their function in transposon silencing, with no detectable function in mammalian somatic tissues. However, PIWI proteins become aberrantly expressed in multiple types of somatic cancers, thus gaining interest in targeted therapy. Despite this, little is known about the regulatory mechanism of PIWI proteins in cancer. Here we report that one of the four PIWI proteins in humans, PIWIL1, is highly expressed in gastric cancer tissues and cell lines. Knocking out PIWIL1 expression (PIWIL1-KO) drastically reduces gastric cancer cell proliferation, migration, metastasis, and tumorigenesis. RNA deep sequencing of gastric cancer cell line SNU-1 reveals that PIWIL1-KO significantly changes the transcriptome, causing the up-regulation of most of its associated transcripts. Surprisingly, few *bona fide* piRNAs exist in gastric cancer cells. Furthermore, abolishing the piRNA-binding activity of PIWIL1 does not affect its oncogenic function. Thus, PIWIL1 function in gastric cancer cells is independent of piRNA. This piRNA-independent regulation involves interaction with the UPF1-mediated nonsense-mediated mRNA decay (NMD) mechanism. Altogether, our findings reveal a novel and piRNA-independent function of PIWIL1 in promoting gastric cancer.

**SIGNIFICANCE:** Precision medicine aims to cure cancer without affecting normal tissues. PIWI proteins provide a promising opportunity for precision medicine because they are normally expressed only in the testis for male fertility but gain expression in diverse types of cancers. Thus, inhibiting *PIWI* expression may stop cancer development (and spermatogenesis) without affecting normal body function. To establish causality between PIWI and cancer, we show here that the expression of PIWIL1, a human PIWI protein, promotes gastric cancer. Surprisingly, this oncogenic function does not require piRNA, the expected partner of PIWI proteins, but involves the nonsense-mediated mRNA decay mechanism. These findings reveal a new function and action mechanism of PIWI proteins in oncogenesis, guiding the identification of PIWI inhibitors to cure cancer.

## INTRODUCTION

Cancer is a malignant disease with a tremendous impact on global health. Surgery, chemotherapy, and radiation therapy are the main types of cancer treatment. Although the overall survival rate of these treatments has improved significantly over the decades, the total therapeutic efficacy is still not ideal, and certain cancers still lack effective therapy. Thus, new treatments, such as immunotherapy, targeted therapies, hormone therapy, and cryoablation, have been developed in recent years for cancer patients (1–4). Among these new types of treatment, targeted therapy has unique strength and potential (2).

Gastric cancer is the fourth most common cancer and the second leading cause of cancer death worldwide (5, 6). Presently, there are only three FDA-approved drugs that target gastric cancer: Trastuzumab (7, 8), Ramucirumab (9–11), and pembrolizumab (12). These drugs have great efficacy in some cancer types such as breast cancer or colon cancer but are much less effective in treating gastric cancer, especially advanced gastric cancer. Therefore, there is a pressing need for new therapy for gastric cancer treatment.

PIWI proteins have emerged as a new opportunity for targeted cancer therapy. These proteins belong to the PAZ-PIWI Domain (PPD) family of RNA-binding proteins, which is composed of the Argonaute and PIWI subfamilies. The PIWI proteins were first discovered for their evolutionarily conserved functions in germline stem cell self-renewal (13, 14). They bind to a class of non-coding small RNAs called PIWI-interacting RNAs (piRNAs) that are generally 24-32 nucleotides in length (15–18). PIWI proteins and piRNAs are mostly expressed in the germline. They form the PIWI-piRNA complex that plays essential roles in germline development, stem cell self-renewal, transposon silencing, and gametogenesis in diverse organisms (19–28). However, the somatic function of PIWI has only been reported in lower eukaryotes (13, 14, 28–30), with the expression and function of PIWIs in mammalian somatic tissues remaining unclear. In mice, complete knockout of all three *PIWI* subfamily genes causes no detectable defects in development and growth (31). In humans, there are four human PIWI genes: *PIWIL1* (*PIWI-Like 1*, a.k.a. *HIWI*), *PIWIL2* (a.k.a. *HILI*), *PIWIL3* (a.k.a. *HIWI3*), and *PIWIL4* (a.k.a. *HIWI2*) (32). *PIWIL1* was first reported to be drastically overexpressed in seminoma, a testicular germ cell tumor (33). Since then, multiple studies have documented the ectopic expression of PIWI proteins in a wide variety of human cancers (34–45). Most of these findings are correlative. Recently, the causative role of *PIWIL1* and *PIWIL4* genes was demonstrated in pancreatic cancer cells and breast cancer cells, respectively (45, 46). In the pancreatic cancer study, the piRNA expression was not detectable, which led the authors to propose that the function of PIWIL1 is piRNA-independent. This proposal is similar to an earlier proposition based on the observation that PIWIL1 did not detectably associate with piRNA in a colon cancer cell line (COLO205) (47). However, some recent studies reported the existence and function of specific piRNAs in cancer, mostly based on correlative evidence such as genetic association studies (48, 49). These contrasting proposals await definitive demonstration of piRNA-independence of PIWI proteins for their function in cancer.

Here we demonstrate a piRNA-independent function of PIWIL1 in gastric cancer. Furthermore, we show that PIWIL1 achieves this function by associating with the UPF1-mediated NMD pathway to regulate mRNA expression. These findings reveal a critical oncogenic mechanism mediated by a PIWI protein that is distinct from the well-established PIWI-piRNA pathway during normal development.

## RESULTS

### *PIWIL1* is highly expressed in gastric cancer patient samples and gastric cancer cell lines

To explore the function of PIWI proteins in gastric cancer, we first examined the expression of all four *PIWI* genes in six different well-studied gastric cancer cell lines, including adhesion and suspension types of cell lines. Only PIWIL1 was highly expressed in all six gastric cancer cell lines at both mRNA and protein levels, above its expression in the normal gastric epithelial cell line GES-1 (Fig. 1A and B; SI Appendix Fig. S1A and B).

**Fig 1.**
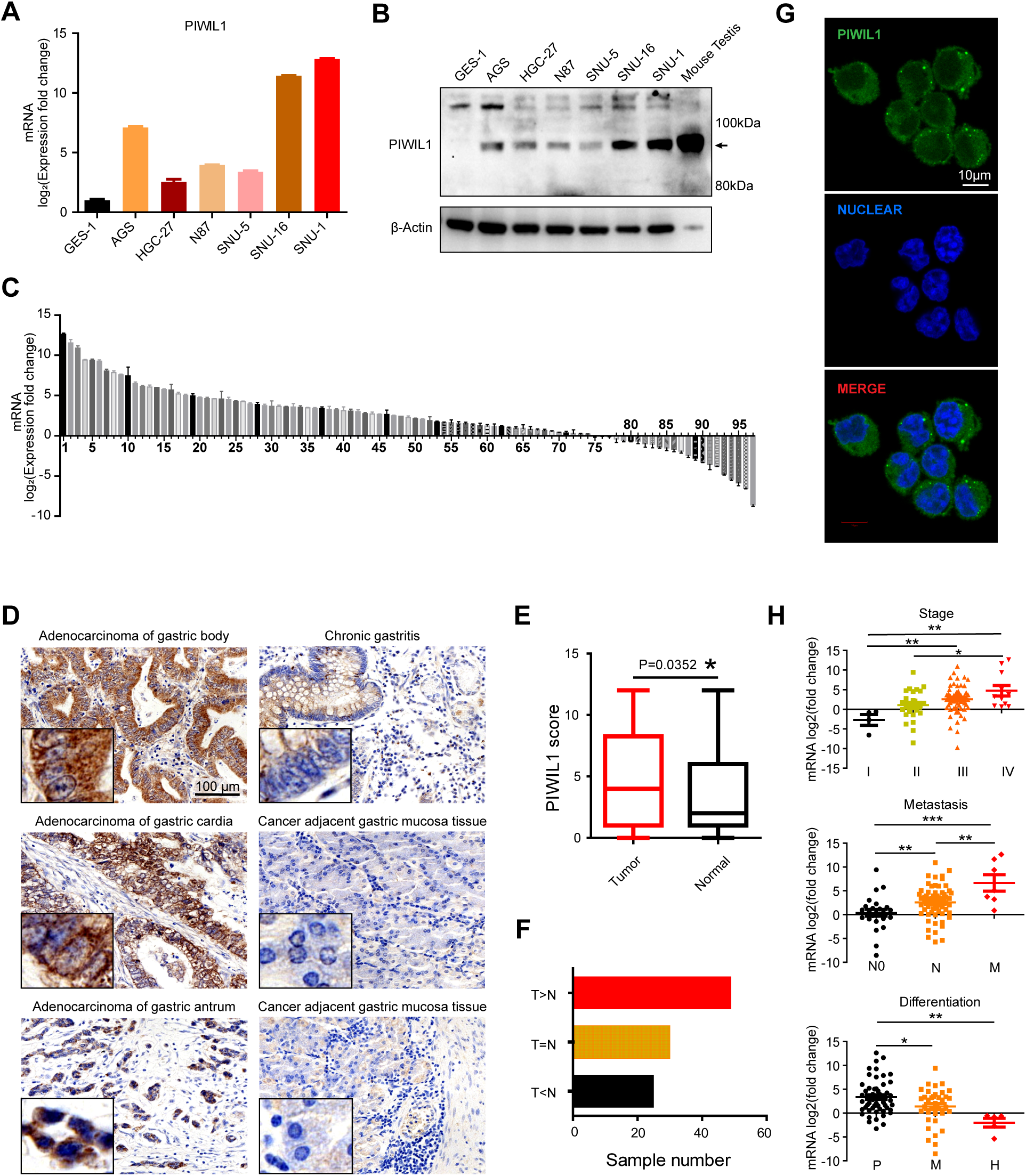
PIWIL1 is highly expressed in gastric cancer samples and gastric cancer cell lines. (A) A bar graph showing quantitative RT-PCR of *PIWIL1* mRNA expression in a normal gastric epithelial cell line GES-1 and six human gastric cancer cell lines. The mRNA level of GES-1 was normalized to 1. β-Actin was used as an internal standard to normalize the level of PIWIL1 from the same sample. (B) A Western blot showing the PIWIL1 protein levels in GES-1 and six human gastric cancer cell lines. β-Actin was used as a loading control. (C) Quantitative RT-PCR examination of the relative PIWIL1 mRNA levels in gastric cancer samples from 97 patients as compared to their paired normal tissues, using β-Actin as an internal control. Bar value represents the difference of PIWIL1 mRNA levels between normal tissue and tumor in log_2_, so that value > 1 and <-1 indicate that PIWIL1 mRNA levels increase and decrease more than two-fold in tumors, respectively. Data were calculated from triplicates. (D) Representative micrographs of PIWIL1 immunohistochemical (IHC) staining of gastric cancer tissue microarrays. (E) A box plot showing IHC scores of PIWIL1 expression in 104 pairs of paired tumor and normal tissues in the gastric cancer tissue microarrays. Each tissue spot on the tissue microarray was scored by stain strength (range from 0 to 3) and the percentage of PIWIL1-positive cells (range from 0 to 4), respectively. The IHC score = stain strength score x percentage score of PIWIL1-positive cells. * p < 0.05, Student’s paired t-Test. (F) PIWIL1-IHC score of each paired tumor and normal tissue spot in gastric cancer tissue microarrays. T>N, T=N, and T<N denote: the number of tissue pairs in which the IHC score in the tumor tissue higher than, equal to, and lower than its paired normal tissue, respectively. The sample number in the T>N group is significantly larger than the sample number in the two other groups. (G) Immunofluorescence staining of PIWIL1 (green) in SNU-1 cells showing the cytoplasmic localization of PIWIL1. The nucleus is stained by DAPI (blue). (H) Bar graphs showing PIWIL1 mRNA levels of 97 paired human gastric cancer samples categorized into different gastric cancer stage, different metastasis state, or different differentiation state. In the upper graph, I, II, III, and IV denote stages 1-4. In the middle graph, N0: no nodes are involved; N: lymph node metastasis including N1, N2, and N3; M: distant metastasis. In the lower graph: P: poor differentiation; M: moderate differentiation; H: high differentiation. Error bars represent standard error. * p < 0.05, ** p < 0.01, *** p < 0.001, Student’s unpaired t-Test.

To further correlate PIWIL1 to gastric cancer, we first conducted a pair-wise comparison of its mRNA expression in gastric cancer samples from 97 patients with their normal gastric tissues. In 63 out of the 97 patients, the level of *PIWIL1* mRNA in the cancer samples was at least two-fold higher than their normal samples (Fig. 1C). Among them, 10 patients had *PIWIL1* mRNA levels that were at least 100-fold higher than their corresponding normal samples. Consistently, the *PIWIL1* mRNA expression data from the Cho gastric dataset of the ONCOMINE database (https://www.oncomine.org) indicate that *PIWIL1*, but not the three other human *PIWI* genes, has higher expression in gastric cancer tissues than normal gastric tissues (SI Appendix Fig. S1C and D). We then examined the PIWIL1 protein expression in a tissue microarray representing 104 gastric cancer patients by immunohistochemical staining. These gastric cancer tissues showed significantly increased levels of the PIWIL1 protein as compared to matched normal tissues (Fig. 1D-F). Most of the PIWIL1 signal was in the cytoplasm (Fig. 1D). Immunofluorescence staining of the gastric cancer cell line SNU-1 also showed the cytoplasmic localization of PIWIL1 (Fig. 1G), consistent with the known role of PIWIL1 as a cytoplasmic protein in mammalian testes (50).

To further correlate the level of PIWIL1 expression to the severity of gastric cancer, we collected and preprocessed 97 gastric cancer patients’ data by extracting four available clinical factors (stage, metastasis, differentiation, and gender). These data indicated that the up-regulation of PIWIL1 mRNA in gastric cancer samples was significantly correlated with both metastasis and the tumor-node-metastasis (TNM) stage but inversely correlated with the degree of differentiation (Fig. 1H; SI Appendix Table S1). The percentage of patients with upregulation of PIWIL1 expression was higher in males than that in females (SI Appendix Table S1). In addition, we assessed the relationship between PIWIL1 expression and clinical outcome of gastric cancer patients. Kaplan-Meier survival analysis showed that when the level of *PIWIL1* mRNA expression was high, patients with poorly differentiated or mixed-classification or 5FU-based adjuvant therapy had a poor prognosis, especially those with poorly differentiated and high TNM stage (SI Appendix Fig. S1E). All these data strongly correlate PIWIL1 with the progression of gastric cancer.

### *PIWIL1* promotes gastric cancer cell proliferation, migration, tumorigenesis, and metastasis

To investigate the effect of aberrant PIWIL1 expression on gastric cancer cells, we chose the gastric cancer cell line SNU-1 as our model. SNU-1 is a poorly differentiated suspension gastric cancer cell line with the highest PIWIL1 expression but low PIWIL2 and PIWIL4 expression among the six gastric cancer cell lines that we examined (Fig. 1A and B; SI Appendix Fig. S1A and B). We deleted a part of the *PIWIL1* gene in SNU-1 cell line by using the CRISPR-Cas9 nickase system (SI Appendix Fig. S2A). This deletion abolished PIWIL1 protein expression (SI Appendix Fig. S2B), and significantly inhibited cell proliferation under normal cell culture condition with 10% fetal bovine serum (FBS), as indicated by two independent knockout (KO) lines (Fig. 2A). Under a malnutritional condition with 5% FBS, the inhibition was even stronger (Fig. 2A). In addition, *PIWIL1*-KO significantly increased the number of cells at G1 phase and decreased the number of cells at S phase, but the number of cells at G2/M phase remained mostly unchanged (Fig. 2B and C). However, *PIWIL1*-KO did not affect apoptosis (SI Appendix Fig. S2C and D). Furthermore, xenografting of SNU-1 cells into nude mice revealed that knocking out *PIWIL1* dramatically retarded the tumor growth of gastric cancer cell SNU-1 *in vivo* (Fig. 2D and E). Immunostaining of sections from these tumors for two cell proliferation markers, PCNA and ki67, revealed that their expression was significantly reduced in the *PIWIL1*-KO tumors (Fig. 2F). Transwell migration assay and tail-vein injection metastasis assay showed that knockout of *PIWIL1* significantly inhibited the gastric cancer cell SNU-1 migration ability *in vitro* (Fig. 2, G and H) and the metastatic potential *in vivo* (Fig. 2I and J). These results indicate that PIWIL1 deficiency severely compromised the SNU-1 gastric cancer cell proliferation, migration, tumorigenesis, and metastasis.

**Fig 2.**
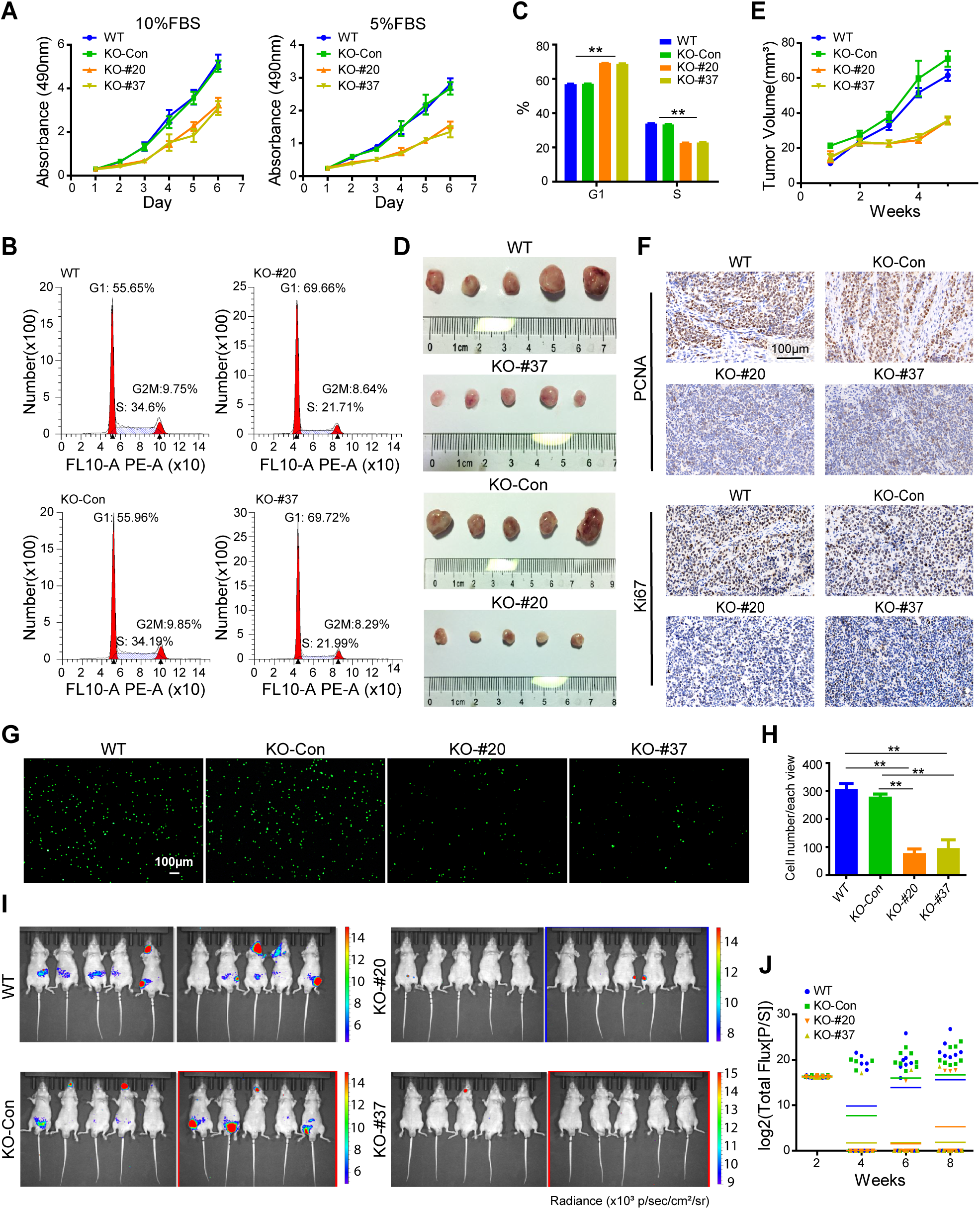
Knockout of *PIWIL1* inhibits SNU-1 cell proliferation, migration, G1-S transition, tumorigenesis, and metastasis. (A) Growth curve of SNU-1 cells with or without PIWIL1-KO in 10%FBS or 5%FBS cell culture medium conditions, analyzed by MTS assay. WT: wildtype SNU-1 cells without transfection of CRISPR-Cas9 plasmids; KO-Con: SNU-1 cells co-transfected with PIWIL1-sgRNA plasmid and Cas9 plasmid but gene-editing did not occur in PIWIL1; KO-#20, KO-#37: two independent SNU-1 cell clones which were co-transfected with PIWIL1-sgRNA plasmid and Cas9 plasmid with gene-editing occurred in PIWIL1 to knockout PIWIL1 protein. Flow cytometry analysis of cell cycle of WT or PIWIL1-KO SNU-1 cells. (B) Flow cytometry analysis of cell cycle of PIWIL1-WT or PIWIL1-KO SNU-1 cells. (C) A bar graph showing the percentage of PIWIL1-WT or PIWIL1-KO SNU-1 cells in G1 and S phase. (D) Representative SNU-1-derived tumors isolated from BNX nude mice 35 days after injection of PIWIL1 WT or KO-control or KO cell clones #20 or #37 SNU-1 cells (5 × 10^6^ cells/mouse). (E) The tumor growth curve of xenograft assay in D. (F) Immunohistochemical staining of cell proliferation markers PCNA and ki67 in the paraffin tumor sections from xenograft-assay in D. (G) Transwell assay of SNU-1 cells with or without knockout of *PIWIL1*. (H) A bar graph showing the number of migrated cell of each well in the transwell assay in G. (I) Representative bioluminescent images of BNX nude mice in either the PIWIL1 WT or KO groups at 8 weeks after injection of SNU-1 cells, depicting the extent of tumor burden. (J) A chart showing quantitative bioluminescent Flux [P/S] of the BNX nude mice in either the PIWIL1 WT or KO groups at 2, 4, 6, and 8 weeks after injection of SNU-1 cells.

To assess whether the *PIWIL1*-KO phenotype we observed in SNU-1 cells reflects a more general function of PIWIL1 in gastric cancer, we performed a similar analysis on another gastric cancer cell line, AGS. After knocking down *PIWIL1* in AGS cell line by three different siRNAs (SI Appendix Fig. S2E), the G1-S transition and migration of AGS cells were all severely affected (SI Appendix Fig. S2F-I). All of the results in this section together demonstrate that PIWIL1 promotes the proliferation, migration, tumorigenesis, and metastasis of gastric cancer cells.

### *PIWIL1* directly targets many RNAs in gastric cancer cells and represses the expression of most of its target RNAs

To explore the molecular mechanism underlying PIWIL1 function in gastric cancer, we carried out total RNA-deep sequencing (RNA-Seq) of wildtype (WT) and *PIWIL1-*KO SNU-1 cells. 1,678 genes are significantly affected in *PIWIL1*-KO cells (adjusted P-value <0.05 & Fold-Change ≥1.5). Weighted gene co-expression network analysis (WGCNA) was used to analyze the RNA-Seq data since changes in the expression of co-expressed gene clusters in *PIWIL1*-knockout cells best reflect the global effect of PIWIL1 on the transcriptome of gastric cancer cells (51, 52). In total, 41 modules were identified via the average linkage hierarchical clustering (Fig. 3A). We then investigated the correlation between PIWIL1 and the co-expressed gene modules. Among the 41 modules, only blue and green modules indicate significantly positive correlations with PIWIL1 expression (p=0.00086 and 0.00014, respectively), whereas the turquoise module shows a significantly negative correlation with PIWIL1 expression (p=6.3e-06) (51) (Fig. 3A and B; SI Appendix Fig. S3A). Especially, genes in blue and turquoise modules are enriched in specific pathways (Fig. 3C; see below). The co-expressed genes in these two modules well represent the difference between WT and *PIWIL1*-KO cells.

**Fig 3.**
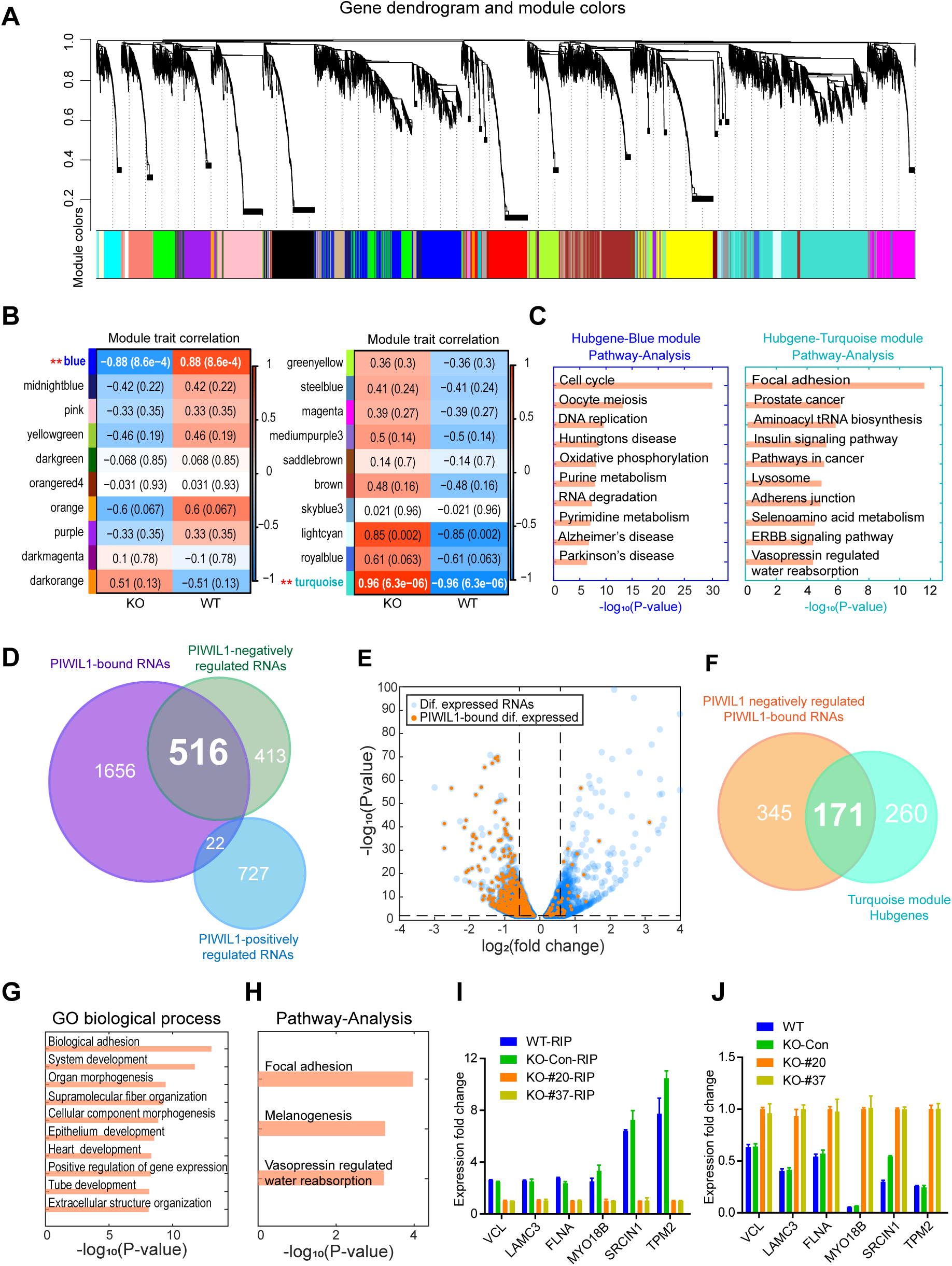
PIWIL1 target RNAs and transcriptomic changes in PIWIL1-KO SNU-1 cells. (A) WGCNA analysis of RNA co-expression modules regulated by PIWIL1. Topological overlap dissimilarity measure is clustered by average linkage hierarchical clustering. Module assignments (using a dynamic hybrid algorithm) are denoted in the color bar (bottom). (B) Heatmap of the correlation between module eigengenes and the trait of with or without PIWIL1 expression. Red color represents a positive correlation between a module and the trait, and blue color represents a negative correlation. Each cell contained the corresponding correlation and *P-value*. (C) KEGG pathway analysis of hub genes of the blue or turquoise module. Any gene with KME ≥0.9 was assigned as a hub gene. GSEA/MSigDB gene sets tool was used for the KEGG pathway analysis. (D) Venn diagram of RNAs positively or negatively regulated by PIWIL1 or bound by PIWIL1. PIWIL1-positively or negatively regulated RNAs are identified by DESeq2 analysis with P<0.05 and fold-change>=1.5 cutoff. PIWIL1-bound RNAs are identified with P<0.05 and fold-change>=1.5 cutoff. (E) Volcano plot of differentially expressed genes (blue dots), including PIWIL1-target genes (orange dots), in PIWIL1-KO SNU-1 cells. Dotted lines represent 1.5-fold-change in expression (vertical lines) and P < 0.01 cutoff (horizontal line). (F) Venn diagram of PIWIL1-bound RNAs negatively regulated by PIWIL1 and RNAs of the turquoise module hub genes with kME ≥0.85 and expression fold-change≥1.5. (G) Gene Ontology (GO) biological process analysis of the 171 PIWIL1-target turquoise hub genes that are negatively regulated by PIWIL1. (H) KEGG pathway analysis of the 171 genes in F and G. (I) Quantitative RIP-PCR validation of PIWIL1-enrichment efficiency of six cell migration-related genes (VCL, LAMC3, FLNA, MYO18B, SRCIN1, and TPM2) from the 171 genes in F-H. (J) Quantitative RT-PCR validation of mRNA expression of the six genes in I.

To further reveal the potential hierarchy of gene regulation in blue and turquoise modules by PIWIL1, we identified 808 and 743 hub genes in the blue and turquoise module, respectively. These hub genes are the most connected in a module and play essential roles in biological processes. The KEGG pathway analysis showed that the cell cycle and DNA replication pathways are highly enriched among the blue module hub genes, whereas the focal adhesion and adherens junction pathways are highly enriched among the turquoise module hub genes (Fig. 3C). The change in the expression of some well-studied oncogenes or tumor suppressor genes in the blue or turquoise module hub genes was validated by quantitative RT-PCR (SI Appendix Fig. S3D). Thus, PIWIL1 appears to promote cell proliferation and cell migration mechanisms but inhibit cell adhesion mechanisms.

To identify the direct target RNAs of PIWIL1 in gastric cancer, we conducted anti-PIWIL1 RNA co-immunoprecipitation (RIP) followed by deep sequencing on SNU-1 cells. This allowed us to identify 2,194 PIWIL1-bound RNAs, among which 538 displayed altered expression in *PIWIL1*-KO cells (Fig. 3D and SI Appendix Fig. S3E). Among these 538 RNAs, 71% were protein-coding mRNAs, 19% were long non-coding RNAs (lncRNAs), and 7% were pseudogene RNAs (SI Appendix Fig. S3F). Remarkably, 516 of the 538 RNAs were up-regulated in *PIWIL1*-KO cells, including 365 mRNA and 101 lncRNAs (Fig. 3D). Only 18 mRNAs enriched in metabolic pathways and 3 lncRNAs were down-regulated in *PIWIL1*-KO cells (SI Appendix Fig. S3G-H). Thus, PIWIL1 negatively regulates most of its target RNAs, regardless of whether they are mRNAs or lncRNAs or even pseudogene RNAs (Fig. 3E and SI Appendix Fig. S3G). Because PIWIL1 is a cytoplasmic protein, it is most likely that PIWIL1 achieves negative regulation by destabilizing its target RNAs instead of repressing their transcription. This post-transcriptional role of PIWIL1 has been demonstrated for mouse PIWIL1 (53). In contrast, 727 indirect target RNAs are positively regulated by PIWIL1, whereas 413 indirect target RNAs are negatively regulated by PIWIL1 (Fig. 3D and SI Appendix Fig. S3G), indicating a preferentially negative regulatory relationship between the PIWIL1-target RNAs and indirect target RNAs.

Among the 516 genes up-regulated in *PIWIL1*-KO cells, 171 were turquoise module hub genes (Fig. 3F) enriched in the adhesion process and the focal adhesion pathway, as revealed by KEGG pathway and Gene Ontology (GO) analyses (Fig. 3G and H). Six of the 171 enriched mRNAs were further confirmed for their binding to, and negative regulation by, PIWIL1 by RIP and qPCR assays (Fig. 3I and J).

Beyond PIWIL1-target RNAs, the expression of another 1,140 RNAs is changed in *PIWIL1*-KO cells, including 221 blue module hub genes that are positively regulated by PIWIL1 and are most highly enriched in the cell cycle pathway (SI Appendix Fig. S3I-J). These findings corroborate the role of PIWIL1 in promoting cell cycle, proliferation, and tumorigenesis. Taken together, our analyses indicate that PIWIL1 achieves its oncogenic function by directly repressing the expression of many genes involved in cell adhesion and indirectly promoting the expression of many genes involved in cell proliferation.

### The PIWIL1 function in gastric cancer cells is piRNA-independent

It has been well demonstrated that PIWI proteins carry out various biological functions by interacting with piRNAs. Our previous work showed that piRNAs derived from transposons and pseudogenes partner with mouse PIWIL1 to degrade many mRNAs and lncRNAs in mouse late spermatocytes (53, 54). We, therefore, investigated whether human PIWIL1 achieves this regulation in gastric cancer via the PIWI-piRNA pathway.

To address this question, we profiled small RNA expression in gastric cancer cells by deep sequencing of the total cellular RNAs and PIWIL1-coimmunoprecipitated RNAs in gastric cancer cell line SNU-1 and used the mouse testis as a positive control. The small RNAs in both WT and *PIWIL1*-KO gastric cancer cells were predominantly miRNAs (22 nucleotides), and there was a barely detectable amount of putative piRNAs (Fig. 4A; Table S2). Furthermore, the total small RNA size profile in WT SNU-1 cells was not different from that in *PIWIL1*-KO SNU-1 samples (Fig. 4A), indicating that knockout of PIWIL1 did not affect the expression of miRNAs, as expected. This is in contrary to mouse testes, in which most of small RNA showed typical piRNA distributions that peak around 30 nucleotides (Fig. 4B) and are not expressed in *PIWIL-*KO mice (53, 54).

**Fig. 4.**
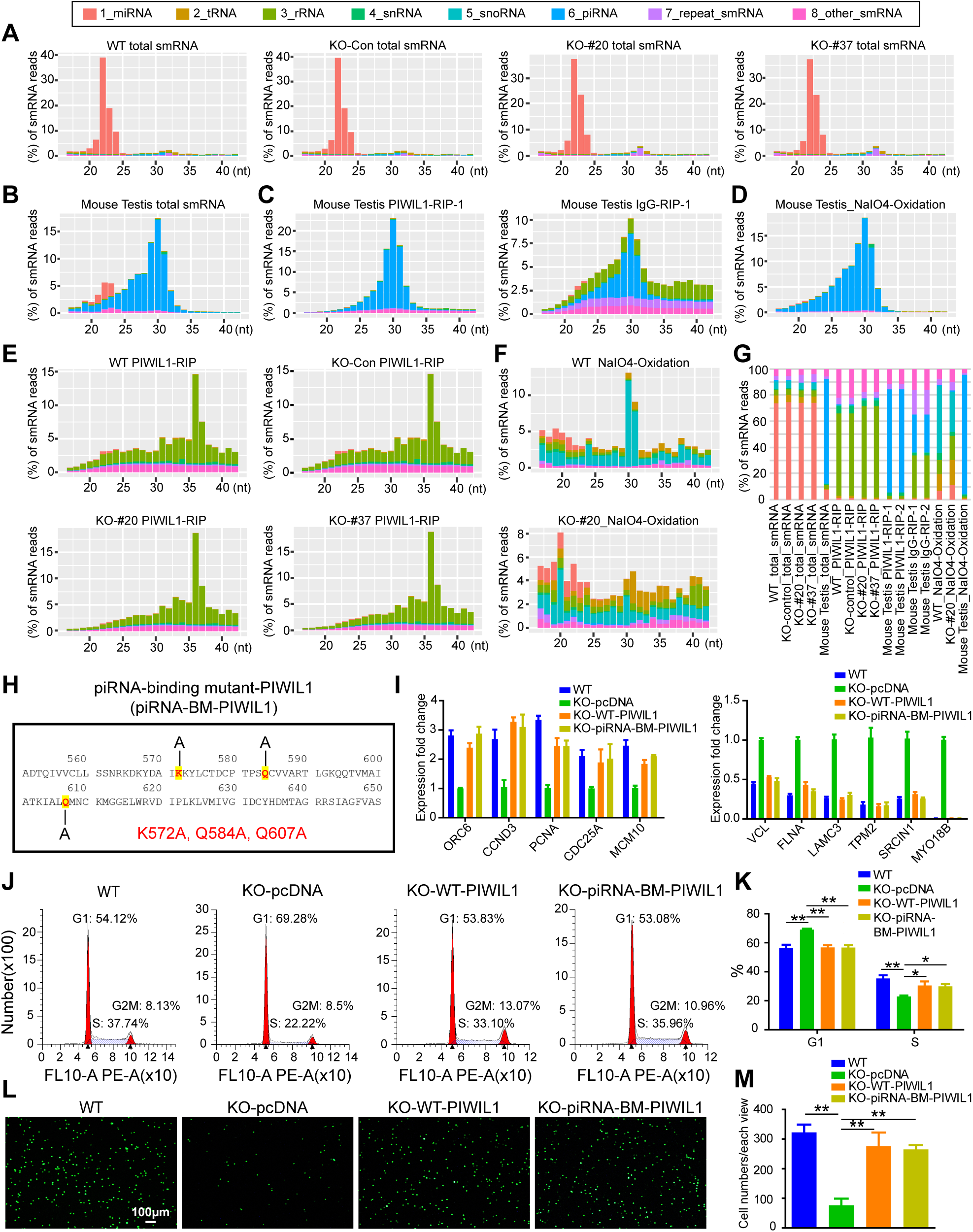
PIWIL1 function in the gastric cancer cell line SNU-1 is piRNA-independent. (A) Size profiles of each class of total small RNAs in WT, PIWIL1-KO-Con, PIWIL1-KO-#20, and PIWIL1-KO-#37 samples. WT: wildtype SNU-1 cells without transfection of CRISPR-Cas9 plasmids; KO-Con: SNU-1 cells co-transfected with PIWIL1-sgRNA plasmid and Cas9 plasmid but gene-editing did not occur in PIWIL1; KO-#20, KO-#37: two independent SNU-1 cell clones which were co-transfected with PIWIL1-sgRNA plasmid and Cas9 plasmid with gene-editing occurred in PIWIL1 to knockout PIWIL1 protein. (B) Size profiles of each class of total small RNAs in mouse testes. (C) Size profiles of each class of small RNAs immunoprecipitated by PIWIL1 antibody in mouse testis, IgG is the IP negative control. (D) Size profiles of each class of the small RNAs in mouse testes after NaIO4 oxidization. (E) Size profiles of each class of small RNAs immunoprecipitated by PIWIL1 antibody in PIWIL1-WT, KO-Con, KO-#20, and KO-#37 samples. (F) Size profiles of each class of small RNAs in PIWIL1-WT and PIWIL1-KO-#20 samples after NaIO4 oxidization. (G) The proportion of each class of small RNAs in Fig. 4 A-F. (H) Mutations in the piRNA-binding mutant PIWIL1 (piRNA-BM-PIWIL1). (I) Left graph: qRT-PCR shows that the downregulation of PIWIL1-target cell cycle mRNAs in PIWIL1-KO cells can be rescued by both WT-PIWIL1 (KO-WT-PIWIL1) and piRNA-binding mutant-PIWIL1 (KO-piRNA-BM-PIWIL1). Right graph: qRT-PCR shows that the upregulation of PIWIL1-target cell migration mRNAs in PIWIL1-KO cells can be rescued by both WT-PIWIL1 (KO-WT-PIWIL1) and piRNA-binding mutant-PIWIL1 (KO-piRNA-BM-PIWIL1). (J) Flow cytometry analysis of cell cycles of WT and PIWIL1-KO SNU-1 cells transfected with the empty vector (KO-pcDNA), as well as PIWIL1-KO cells transfected with wildtype or piRNA-binding mutant PIWIL1-expressing plasmids, denoted as KO-WT-PIWIL1 and KO-piRNA-BM-PIWIL1, respectively. (K) The bar graph shows the percentage of the G1 phase and S phase of WT-PIWIL1, KO-pcDNA, KO-WT-PIWIL1, and KO-piRNA-BM-PIWIL1 SNU-1 cells in J. (L) Transwell assay of WT-PIWIL1, KO-pcDNA, KO-WT-PIWIL1, and KO-piRNA-BM-PIWIL1 SNU-1 cells. (M) The bar graph shows the number of migrated cells in each well in the Transwell assay in L.

To further search for the possible existence of low abundant piRNAs in WT and *PIWIL1*-KO SNU-1 cells that might not have been detected among other small RNAs, we used an anti-PIWIL1 antibody to enrich PIWIL1-bound piRNAs and NaIO_4_ oxidation treatment to remove the 2’O methylation at the 3’end of piRNAs to facilitate piRNA cloning (55, 56). Mouse testes were used as a positive control. Expectedly, piRNAs were enriched in the mouse testicular small RNA preparations after both PIWIL1 antibody pull-down and NaIO_4_-treatment but not in the IgG pull-down control (Fig. 4B-D; SI Appendix Fig. S4A), indicating the validity of piRNA enrichment by these two methods. However, small RNA preparations from SNU-1 cells showed no piRNA enrichment but the spurious distribution of small RNA categories mainly represented by degraded rRNA fragments in RIP libraries (Fig. 4E; SI Appendix Tables S2-3) and snoRNAs in NaIO_4_-treated libraries (Fig. 4F). These lines of evidence support that very few *bona fide* piRNAs, if any, exist in the SNU-1 gastric cancer cell line. In addition, the expression of several key components for piRNA biogenesis was barely detected (SI Appendix Fig. S4B), further indicating the absence of a functional piRNA biogenesis machinery in SNU-1 cells.

To further demonstrate that PIWIL1 functions independent of piRNAs in SNU-1 cells, we investigated whether mutating piRNA-binding residues in PIWIL1 will compromise its function. We generated a transfection construct containing a piRNA-binding mutant *PIWIL1* by introducing K572A, Q584A, and Q607A triple mutations into a WT *PIWIL1* cDNA (57, 58) (Fig. 4H). The mutant *PIWIL1* was introduced into *PIWIL1*-KO SNU-1 cells by transfection. The WT *PIWIL1* gene and the empty vector were used as a positive and negative control, respectively. The mutant *PIWIL1* fully restored the regulation of the PIWIL1 target RNAs in *PIWIL1*-KO SNU-1 gastric cancer cells, as did the WT *PIWIL1* transgene (Fig. 4I). Notably, just like WT PIWIL1, the mutant PIWIL1 blocked the down-regulation of *PIWIL1*-KO-induced cell cycle-associated mRNAs (*CCND3, ORC6, PCNA, CDC25A*, and *MCM10*) and the up-regulation of *PIWIL1*-KO-induced cell migration-associated mRNAs (*VCL, FLNA, LAMC3, TPM2, SRCIN1*, and *MYO18B*)(59–66). Moreover, the mutant-PIWIL1 rescued the cell cycle arrest and cell migration defects in *PIWIL1*-KO SNU-1 cells comparable to WT PIWIL1 (Fig. 4J-M). Together, these lines of evidence indicate that the function of PIWIL1 in gastric cancer cells is independent of piRNA.

### The piRNA-independent regulation of PIWIL1 involves the NMD mechanism

To investigate how PIWIL1 carries out piRNA-independent regulation in gastric cancer cells, we performed PIWIL1-coimmunoprecipitation followed by mass spectrometry to identify proteins that interact with PIWIL1 in SNU-1 cells. The PIWIL1 antibody effectively pulled down PIWIL1 protein itself and UPF1 (Up-Frameshift-1), a core factor of nonsense-mediated decay (NMD) of mRNAs (67, 68)(Fig. 5A). This indicates that PIWIL1 might negatively regulate its direct target RNAs by interacting with the UPF1-mediated NMD complex.

**Fig. 5.**
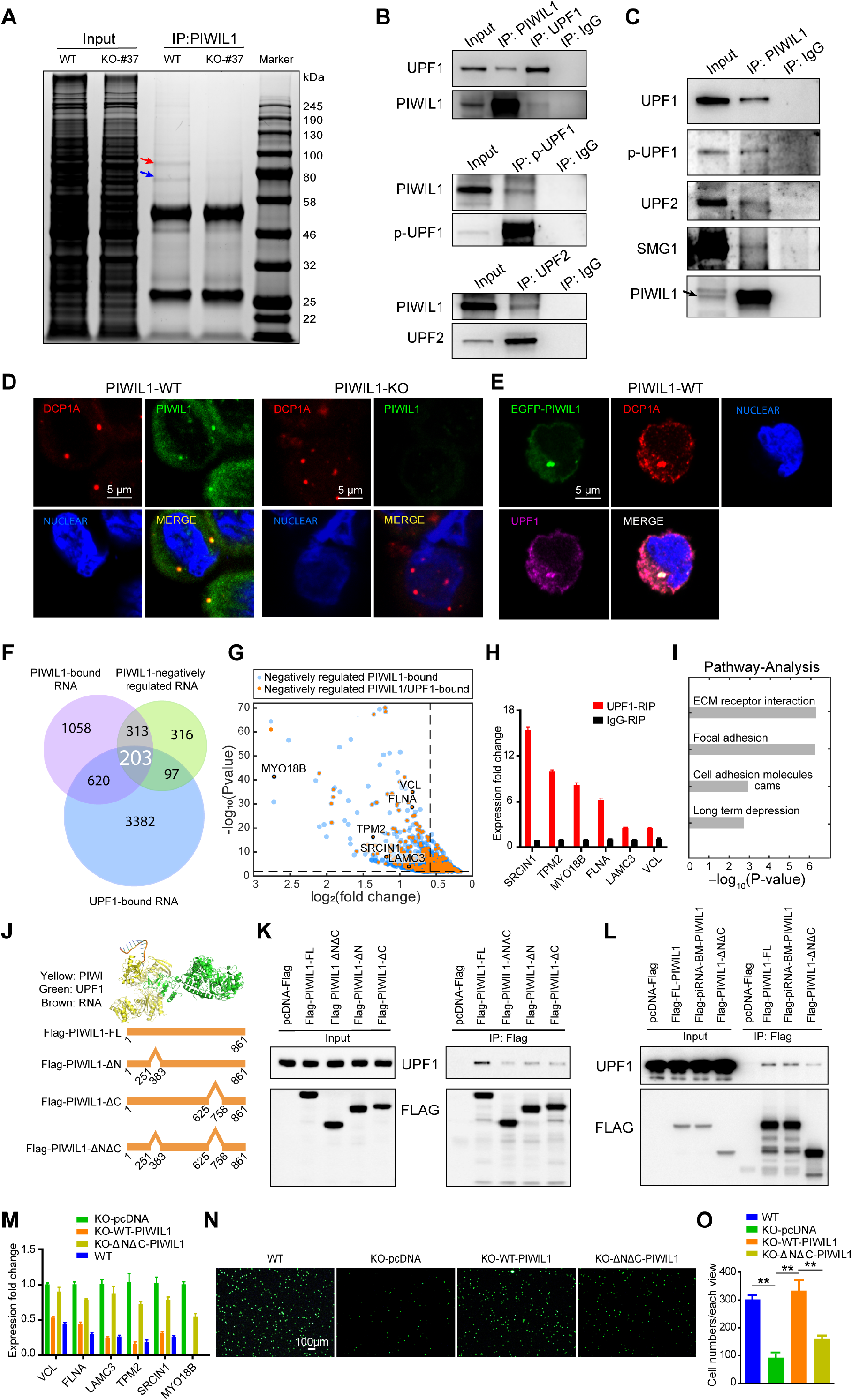
PIWIL1 cooperates with UPF1 to negatively regulate PIWIL1-bound RNAs in a piRNA-independent manner. (A) A Coomassie blue staining protein gel of PIWIL1-co-immunoprecipitation, Mass spectrometry of this protein gel identifies PIWIL1 (blue arrow) as well as UPF1 (red arrow). (B) Western blotting showing reciprocal co-immunoprecipitated between PIWIL1 and UPF1 (total and phosphorylated form, p-UPF1) and between PIWIL1 and UPF2. (C) Western blotting shows that PIWIL1 co-immunoprecipitates with NMD complex core proteins UPF1, phosphorylated UPF1 (p-UPF1), UPF2, and SMG1. (D) Immunofluorescence staining of PIWIL1 (green) and DCP1A (red, a P body marker) in wildtype and PIWIL_KO SNU-1 cells. (E) Immunofluorescence staining of PIWIL1 (green), DCP1A (red), and UPF1 (fuchsia) in SNU-1 cells, which shows PIWIL1 co-localized with UPF1 in the P body. (F) Venn diagram of PIWIL1-bound RNAs, UPF1-bound RNAs, and PIWIL1-negatively regulated RNAs, with P<0.05 cutoff and fold-change>=1.5 cutoff. 203 PIWIL1-negatively regulated RNAs are targeted by both PIWIL1 and UPF1. (G) Volcano plot of PIWIL1 negatively regulated direct-targets (blue dots) in SNU-1 cells. Orange dots are PIWIL1 negatively regulated direct-targets, which are also targeted by UPF1. Dots demarcated by black outlines are six cancer cell migration-related turquoise module hub genes (VCL, FLNA, LAMC3, SRCIN1, TPM2, and MYO18B). Dotted lines of the Volcano plot represent 1.5-fold-change in expression (vertical lines) and P < 0.01 cutoff (horizontal line). (H) Quantitative RIP-PCR confirmed that the six cancer cell migration-related turquoise module hub genes are bound by UPF1. (I) KEGG pathway analysis of 203 PIWIL1-negatively regulated RNAs that are targeted by both PIWIL1 and UPF1. (J) Docking model of PIWIL1 protein structure (yellow), UPF1 protein structure (green), and RNA (orange). Schematic of full length-PIWIL1 and UPF1-interacting domain mutants-PIWIL1. (K) Co-immunoprecipitation mapping of the UPF1-interacting domain of PIWIL1. pcDNA-Flag: empty vector, Flag-PIWIL1-FL: full-length PIWIL1, Flag-PIWIL1-ΔN: Flag-tagged PIWIL1 without 251-383 amino acids, Flag-PIWIL1-ΔC: Flag-tagged PIWIL1 without 624-758 amino acids, Flag-PIWIL1-ΔNΔC: Flag-tagged PIWIL1 without both 251-383 and 624-758 amino acids. (L) Western blotting of co-immunoprecipitation showing that piRNA-binding mutant PIWIL1 (Flag-piRNA-BM-PIWIL1) interacts with UPF1 in the same way as WT PIWIL1 (Flag-PIWIL1-FL). (M) Quantitative RT-PCR showing that the upregulation of PIWIL1-UPF1 co-targeted mRNAs in PIWIL1-KO cells can be rescued by WT-PIWIL1 but not the mutant PIWIL1 lacking the UPF1-interacting domain (KO-ΔNΔC-PIWIL1). (N) Transwell assays showing that the inhibition of cell migration in PIWIL1-KO cells can be rescued by WT-PIWIL1 but not the mutant PIWIL1 lacking the UPF1-interacting domain (KO-ΔNΔC-PIWIL1). (O) The bar graph shows the numbers of migrated cells of each view in the Transwell assay.

To investigate this possibility, we conducted co-immunoprecipitation assays between PIWIL1 and two core factors of the NMD machinery, UPF1, and UPF2. We found that PIWIL1 interacted with both UPF1 and UPF2. Since UPF1 is phosphorylated by SMG1 and then recruits various decay effectors of the NMD degradation machinery to target RNAs, we further explored whether PIWIL1 interacts with SMG1 and/or phosphorylated-UPF1 (p-UPF1) by co-immunoprecipitation of PIWIL1 and SMG1 as well as PIWIL1 and p-UPF1 using SNU-1 cellular extracts. PIWIL1 interacted with SMG1 and p-UPF1 in addition to UPF1 and UPF2 (Fig. 5B and C). Moreover, the interactions between PIWIL1 and the NMD pathway core factors were also observed in another gastric cancer cell line AGS (SI Appendix Fig. S5A), confirming that PIWIL1 interacts with the NMD machinery in gastric cancer cells.

To confirm that PIWIL1 is involved in the NMD pathway, we conducted immunofluorescence microscopy to determine whether PIWIL1 and UPF1 co-localize in SNU-1 cells. It was well documented that UPF1 triggers mRNA decay in P-bodies, which are large cytoplasmic granules replete with proteins involved in general mRNA decay and related processes (69–71). Indeed, PIWIL1 is co-localized with UPF-1 in P-bodies in WT SNU-1 cells, but PIWIL1 signal was not detected in *PIWIL1*-KO SNU-1 cells (Fig. 5D and E). In addition, PIWIL1 is localized in P-bodies in AGS cells (SI Appendix Fig. S5B). These observations indicate that PIWIL1 interact with the NMD pathway components in the P body, and further validate the PIWIL-UPF1 interaction.

To identify RNA co-targets of PIWIL1 and the NMD pathway, we compared PIWIL1-RIP-seq data with the UPF1 RIP-seq data of SNU-1 cells and identified 203 mRNAs that are negatively regulated by PIWIL1 and bound by both PIWIL1 and UPF1 (Fig. 5F and G). The binding of these PIWIL1-target RNAs by UPF1 was confirmed by UPF1 RIP-qPCR analysis of six representative target hub mRNAs (*VCL*, *FLNA*, *LAMC3*, *SRCIN1*, *TPM2*, and *MYO18B* mRNAs, Fig. 5H). KEGG pathway analysis of these 203 co-target RNAs shows that they are highly enriched in extracellular Matrix (ECM) receptor interaction, focal adhesion, and cell adhesion proteins (Fig. 5I and SI Appendix Fig. S5C), indicating that PIWIL1 appears to coordinate with UPF1 to regulate these co-targeting mRNAs that may play a negative role in gastric cancer cell migration.

We then mapped specific domains responsible for PIWIL1 interaction with UPF1 and investigated whether such a domain is required for the regulation of PIWIL1-UPF1 co-targeted genes. We developed a UPF1 and PIWIL1 protein structure-docking model to guide domain mapping (72–74) (Fig. 5I). Deletion of 251-383 amino acid residues and 625-758 residues of PIWIL1 abolished its interaction with UPF1, but the piRNA-binding mutant PIWIL1 interacted with UPF1 in the same way as WT PIWIL1 (Fig. 5J-L). Moreover, the deletion mutant PIWIL1 failed to prevent the up-regulation of all six PIWIL1 and UPF1 co-targeted hub genes that we examined in *PIWIL1*-KO SNU-1 cells (Fig. 5M). This indicates the importance of PIWIL1-UPF1 interaction for PIWIL1 regulation towards its target genes. In contrast, the piRNA-binding mutant PIWIL1 acted like WT PIWIL1 in preventing the up-regulation of these hub genes caused by *PIWIL1*-KO (Fig. 4I). In addition, the deletion mutant PIWIL1 failed to rescue the decreased cell migration of *PIWIL1*-KO cells (Fig. 5N-O), whereas the piRNA-binding mutant-PIWIL1 fully rescued the cell migration defects of *PIWIL1*-KO as did WT-PIWIL1 (Fig. 4L and M). All these observations indicate that interaction with UPF1 is essential for PIWIL1 function in SNU-1 cells.

Finally, we directly assessed the expression of UPF1 in gastric cancer patient samples. In our 97 paired gastric cancer patient samples, *UPF1* mRNA was overexpressed in 74 patients as compared to their corresponding normal samples (SI Appendix Fig. S6A). Consistent with our results, the GEPIA database (http://gepia.cancer-pku.cn/) indicates that *UPF1* mRNA expression in gastric cancer samples is higher than that in normal tissue samples (SI Appendix Fig. S6B). Finally, the expression of *PIWIL1* mRNA is significantly correlated with the expression of *UPF1* mRNA in our 97 paired clinical patient samples (Pearson-*p*=0.0458; SI Appendix Fig. S6C). Collectively, these results indicate that UPF1 is a crucial partner of PIWIL1 in gastric cancer and that PIWIL1 acts through the NMD mechanism in a piRNA-independent fashion to negatively regulate many of its target RNAs in gastric cancer cells.

## DISCUSSION

Although environmental factors such as high-fat diet and *H. pylori* are better known to cause gastric cancer, genetic aberrations have been increasingly recognized to play an important role. PIWI proteins have been reported to be ectopically expressed in diverse types of cancer, but their role in gastric cancer or most other types of cancer has not been established. Here we report that PIWIL1 is aberrantly expressed in both gastric cancer tissues and cell lines, which promotes the progression of gastric cancer (Fig. 1). Inhibiting the aberrant PIWIL1 expression reduces cancer cell proliferation, migration, tumorigenesis, and metastasis (Fig. 2 and SI Appendix Fig. S2). These findings reveal a novel function of PIWIL1 in gastric cancer and identify PIWIL1 as a potential target for gastric cancer precision therapy.

It has been well documented that PIWIL1 regulates the expression of a range of genes at transcriptional or post-transcriptional levels in the germline by forming a complex with piRNAs. To our knowledge, this study represents the first transcriptomic analysis of genes regulated by a PIWI protein in gastric cancer cells. Our work identified 538 RNAs bound and directly regulated by PIWIL1. PIWIL1 regulates most of them negatively (Fig. 3D-F). These RNAs are enriched in migration-related tumor suppressor genes (Fig. 3A-C; SI Appendix Fig. S3A-C). In addition, PIWIL1 positively but indirectly regulates many cell cycle-related oncogenes (Fig. 3A-C). These findings indicate an overall hierarchy of negative regulation by PIWIL1 to its direct targets, such as tumor repressor. These direct targets, in turn, preferentially exert negative regulation towards the indirect targets of PIWILI, including genes that promote cell proliferation and oncogenesis.

A major surprising finding from this study is the piRNA-independent nature of PIWIL1 function. This is in contrast to the well-established piRNA-dependent function of PIWI proteins both in the germline and in the soma. The gastric cancer cells express very few *bona fide* piRNAs if any. Although there was a report of the existence of piRNAs in diverse somatic tissues in the mouse and macaque (75) and several reports of piRNA expression in cancers (76–79), tRNA fragments in sperm derived from tRNA-Gly or tRNA-Glu share nearly identical sequences to some of the annotated piRNAs (80–82), so the authenticity and function of some of the reported somatic piRNAs deserve further investigation. Our finding is consistent with two latest studies on a colon cancer cell line (COLO205) and pancreatic ductal adenocarcinoma cells, respectively (46, 47). However, unlike the COLO205 cell line in which the knockout of *PIWIL1* is inconsequential to the cancer cell transcriptome, we detected significant changes in the transcriptome. Furthermore, we identified a piRNA-independent function of PIWIL1 via interacting with UPF1 and other core components of the NMD mechanism to negatively regulate the expression of its direct target mRNAs (Fig.5). Specifically, both WT and piRNA-binding-deficient PIWIL1 interact with UPF1 equally effectively with similar regulatory effects and biological impact. Such interaction is essential for PIWIL1 function. The difference between our findings and the COLO205 study could reflect the different requirements of PIWIL1 in different types of cancers. This possibility can be tested by investigating whether PIWIL1 has any oncogenic function in COLO205 cells. In any case, our findings identified a novel mechanism of PIWIL1 action by interacting with the NMD machinery in a piRNA-independent manner.

The NMD machinery executes mRNA degradation through at least two pathways: 3ʹ UTR exon junction complex (EJC)-dependent NMD pathway that degrades mRNA containing premature stop codons and 3ʹ UTR EJC-independent NMD pathway that degrades normal mRNAs (83, 84). Both pathways involve UPF1 and SMG1 kinase, and often UPF2 and UPF3 (84, 85). In addition, in both pathways, UPF1 and SMG1 are recruited to the target mRNA, where SMG1 phosphorylates UPF. This phosphorylation activates UPF1 to further recruit downstream components to degrade the target mRNA. Given that the PIWIL1-UPF1 co-target mRNAs are presumably normal, it is likely that PIWIL1 moderates the activity of the NMD machinery through the EJC-independent pathway and at least in part through interacting with the phosphorylated form of UPF1. The future systematic analysis will test this possibility and precisely determine the step(s) of the NMD pathway at which PIWIL1 exerts its regulatory function.

All findings in this report converge into an overall model on how PIWIL1 might promote gastric cancer (Fig. 6): PIWIL1 forms a complex with UPF1, UPF2, SMG1, and other components of the NMD machinery in the P body, possibly through the EJC-independent NMD pathway, to degrade its target mRNAs and lncRNAs, especially tumor suppressors mRNAs, This compromises the overall negative regulation of PIWIL1-target genes towards many indirect target mRNAs, including those for cell proliferation and oncogenesis, thus allowing these mRNAs to be well expressed to promote cancer development. Meanwhile, the PIWIL1-NMD pathway also directly degrades some mRNAs for proteins that inhibit migration, such as cell adhesion molecules, which also promotes oncogenic development. Overall, our findings start to reveal a piRNA-independent function of PIWI proteins in partnership with the NMD mechanism, a novel gastric cancer-promoting mechanism, and a potential therapeutic target for precision therapy of gastric cancer and possibly other types of cancers.

**Fig. 6.**
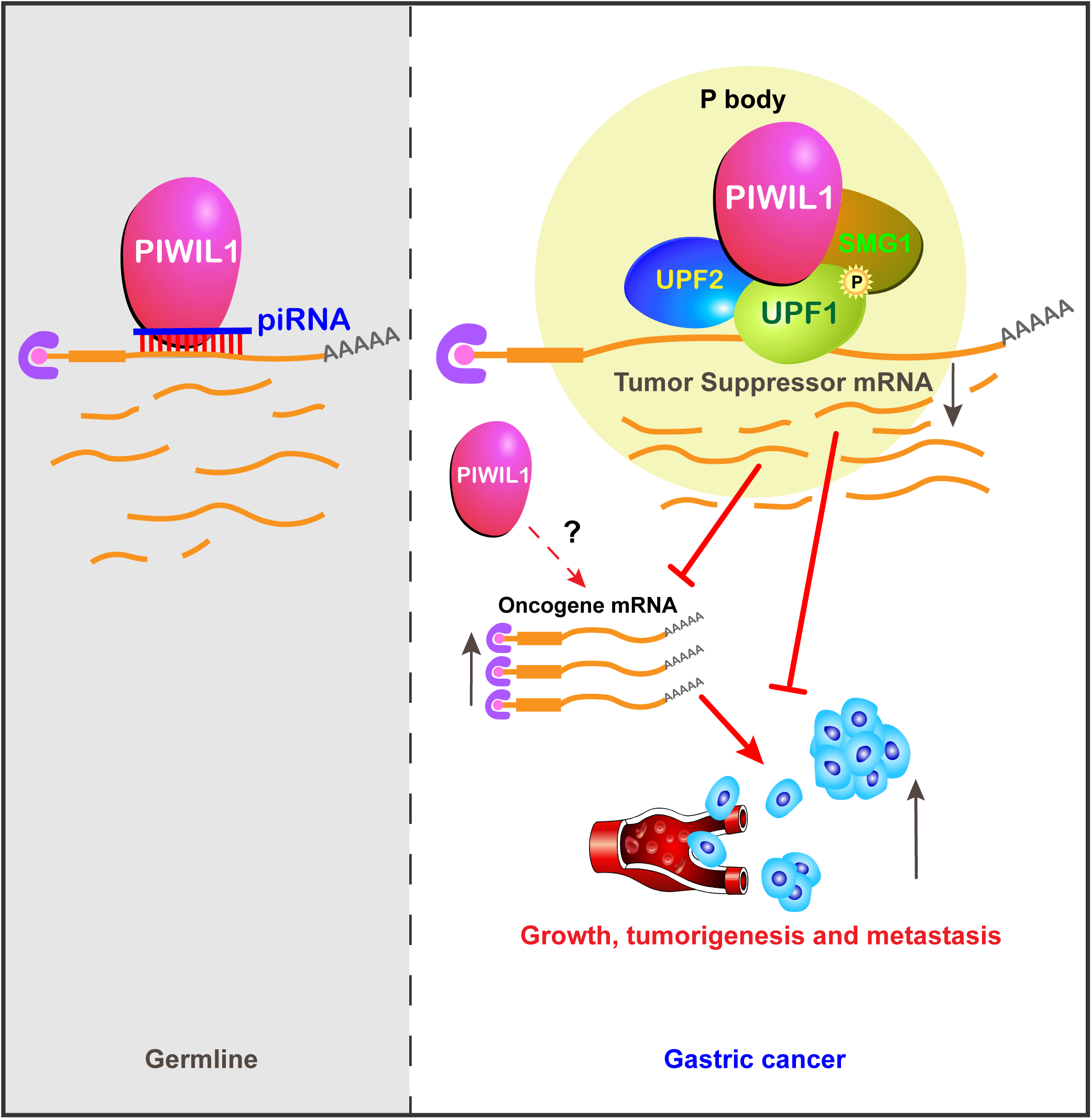
A piRNA-independent mechanism of PIWIL1 in regulating mRNA turnover in gastric cancer cells (right panel), in contrast to the canonical PIWIL1 action mechanism in the germline cytoplasm in which the targeting of PIWIL1 towards a mRNA is guided by the piRNA complementarity with the mRNA. For details, see the main text.

## MATERIALS AND METHODS

### Cell culture and clinical samples

SNU-5 cells were cultured in IMDM medium (ThermoFisher Scientific, 31980030) supplemented with 20% fetal bovine serum. SNU-1, SNU-16, N87, and HGC-27 cells were cultured in RPMI 1640 medium (ThermoFisher Scientific, 61870036) supplemented with 10% fetal bovine serum. AGS cells were cultured in ATCC-formulated F-12K Medium (ATCC, 30-2004) supplemented with 10% fetal bovine serum. GES-1 cells were cultured in DMEM medium (ThermoFisher Scientific, 11995065) supplemented with 10% fetal bovine medium. All these cell lines were incubated at 37 °C with 5% CO2.

Ninety-seven pairs of clinical samples were purchased from the tissue bank of the Institute of Health Sciences, Chinese Academy of Sciences. The local ethics committee approved the study, and the regulations of this committee were followed.

### RNA extraction and quantitative RT-PCR

Total RNA was isolated using TRIzol (Invitrogen) according to the protocol of the manufacturer. For reverse transcription, we used the ABI High Capacity cDNA Reverse Transcription kit (Life Technologies, 4368814). Quantitative RT-PCR reactions were performed according to the protocol of the Bio-Rad real-time PCR system (iQTM SYBR Green Supermix and CFX96TM real-time system). Primers of β-Actin were designed as a real-time PCR control. Quantitative PCR primers are listed in supplemental Table S4.

### *PIWIL1* and *PIWIL1* mutant cDNA cloning

Total RNA was used for cDNA synthesis by SuperScript_III reverse transcriptase (Invitrogen, 18080044) according to the protocol of the manufacturer. The cDNA was used as a template for amplification by Phusion high-fidelity DNA polymerase (New England Biolabs, M0530L) in PCR and PIWIL1 was cloned into the pcDNA3.1-3xFlag.

The cloning of the piRNA-binding mutant *PIWIL1* gene was carried out by the KOD-Plus-Mutagenesis kit (TOYOBO, SMK-101) with pcDNA3.1-3x*Flag-PIWIL1* cDNA as a template, according to the protocol of the manufacturer. PCR primers are listed in supplemental TableS5. The cloning of PIWIL1-domain mutants was carried out by the Phanta Max Super-Fidelity DNA Polymerase kit (Vazyme, P505-d1) and Mut Express MultiS Fast Mutagenesis Kit V2 (Vazyme, C215-01), according to the protocol of the manufacturer. Primer (2-250)/(2-624)-F’ and Primer (2-250)-R’ were used to get fragment (2-250); Primer (384-861)/(384-624)-F’ and Primer (384-861)/(759-861)-R’ were used to get fragment (384-861); Primer (2-250)/(2-624)-F’ and Primer (2-624)/(384-624)-R’ were used to get fragment (2-624); Primer (759-861)-F’ and Primer (384-861)/(759-861)-R’ were used to get fragment (759-861); Primer (384-861)/(384-624)-F’ and Primer (2-624)/(384-624)-R’ were used to get fragment (384-624). The *flag-PIWIL1-ΔN* mutant cDNA (deletion of 251-383) was constructed by connecting fragment (2-250) and fragment (384-861). The *flag-PIWIL1-ΔC* mutant cDNA (deletion of 625-758) was built by connecting fragments (2-624) and fragment (759-861). The *flag-PIWIL1-ΔNΔC* mutant cDNA (deletion of 251-383 and 625-758) was built by connecting fragment (2-250) to fragment (384-624) and fragment (759-861). PCR primers are listed in supplemental TableS5.

### Western Blotting Analysis

Total proteins were extracted by lysis buffer [20mM Tris-HCl pH7.4, 150mM NaCl, 1 % (v/v) IGEPAL^®^ CA-630 (Millipore SIGMA, Cat# I8896), 1mM EDTA, 0.5 mM DTT, cOmplete™ EDTA-free Protease Inhibitor Cocktail Tablets (MERCK, Cat# 4693132001), and PhosStop Tablets (MERCK, Cat# 4906837001)]. Protein samples were diluted (3:1) with a 4x Laemmli sample buffer (Biorad, Cat# 1610747) and heated at 98 °C for 5 min. Proteins were resolved by the TGX Fast Cast acrylamide kit, 7.5% or 10% (Bio-Rad, 1610173TA) at 120 V, and electro-transferred to a PVDF membrane (Merck/Millipore, IPVH00010) at 0.3A for 1.5 h. The membrane was blocked with 5% DifcoTM skim milk (BD Biosciences, 232100) at room temperature for 2 h, which was diluted with TBS (Bio-Rad, 1706435) supplemented with 0.1% Tween 20 (Santa Cruz Biotechnology, sc-29113). PIWIL1 antibody (Abcam, ab181056) was used at 1:1000 dilutions. PIWIL2 antibody (Abcam, ab138742) was used at 1:100 dilution. PIWIL4 antibody (Abcam, ab111714) was used at 1:1000 dilution.

### Cell Proliferation and Apoptosis Assays

Cell proliferation was determined using CellTiter 96_AQueous MTS (3-(4,5-dimethylthiazol-2-yr)-5-(3-carboxymethoxyphenyl)-2-(4-sulfophenyl)-2H tetrazolium, inner salt) reagent powder (Promega, G1111) according to the instructions of the manufacturer. Absorbance at 490-nm wavelength was read using EnSpire_multimode plate readers (PerkinElmer Life Sciences).

Cell apoptosis was determined using FITC Annexin V apoptosis detection kit I (BD Biosciences, 556547) according to the protocol of the manufacturer and analyzed.

### Transwell migration assay

The transwell assay was done using Corning FluoroBlokTM cell culture inserts (Falcon, 351152) according to the protocol of the manufacturer. 1×10^5^ cells were seeded in each well. After three hours, cells were stained and photographed.

### *In vivo* tumorigenicity assay

Five-week-old male nude mice were housed under standard conditions. SNU-1 gastric cancer cells were harvested and washed with PBS and suspended in RPMI 1640 medium without serum. 5 × 10^6^ of these cells were injected subcutaneously into the right flank of nude mice. Tumor growth was measured every 7 days, and tumor volume was estimated as length × width × height × 0.5236. Tumors were harvested from ether-anesthetized mice. All procedures were in agreement with the Guide for the Care and Use of Laboratory Animals (NIH publication nos. 80-23, revised 1996), and approved by the Animal Care and use Committee, ShanghaiTech University.

### *In Vivo* Metastasis Assay

The SNU-1 gastric cancer cell line was labeled with luciferase-expressing lentivirus containing an independent open reading frame of GFP. Luciferase expression was determined by using luciferin (Xenogen, Alameda, CA) and an *in vivo* imaging system (Xenogen). The luciferase-expressing SNU-1 of PIWIL1-WT or PIWIL1-KO cells (1 × 10^6^ in 100 μl PBS) were injected through the tail vein, after which the overall health condition and bodyweight of the mice were monitored. The metastatic lesions were monitored every other week. An aqueous solution of luciferin (150 mg/kg intraperitoneally) was injected 10 min before imaging, and then the mice were anesthetized with Forane (Abbott). The mice were placed into a light-tight chamber of the CCD camera system (Xenogen), and the photons emitted from the luciferase-expressing cells within the animal were quantified for 1 min using the software program Living Image (Xenogen) as an overlay on Igor (Wavemetrics, Seattle, WA).). All procedures were in agreement with the Guide for the Care and Use of Laboratory Animals (NIH publication nos. 80-23, revised 1996) and approved by the Animal Care and use Committee, ShanghaiTech University.

### Immunohistochemistry

Human gastric cancer tissue chip (Cat# HStm-Can090PT-01) was purchased from Shanghai Superchips Company. Mouse tumors for immunohistochemistry were harvested from xenograft mice, cut into 10-micrometer-thick consecutive sections, and mounted on glass slides. After deparaffinized, rehydrating, antigen retrieval, and blocking endogenous peroxidases, the sections or tissue chip were washed three times in PBS for 5 mins each and blocked for 1 h in 0.01 mol/L PBS supplemented with 0.3% Triton X-100 and 5% normal goat serum, followed by addition of anti-PIWIL1 antibody (Atlas antibodies, HPA018798, 1:1000 dilution), or anti-PCNA antibody (Servicebio, GB11010-1, 1:1000 dilution), or anti-Ki67 antibody (Servicebio, GB13030-2, 1:300 dilution) at 4°C overnight. After brief washes in 0.01 mol/L PBS, sections were exposed for 2 hours to 0.01 mol/L PBS containing horseradish peroxidase-conjugated rabbit anti-goat immunoglobulin G (1:500), followed by development with 0.003% H2O2 and 0.03% 3,3′-diaminobenzidine in Tris-HCl (pH 7.6). Immunohistochemistry for each sample was performed at least three separate times, and all sections were counterstained with hematoxylin.

### Immunofluorescence Microscopy

2×10^5^ cells were seeded on a coverslip (Fisher, 12-545-83) in a 24-well plate. After 24 h, cells were washed three times in PBS and then fixed in 4% formaldehyde (paraformaldehyde powder, 95%, 158127-2.5KG, Sigma) at room temperature for 15 min. The fixed cells were washed in ice-cold PBST (1% Tween 20 in PBS) three times (5 min each wash), blocked in 3% BSA in PBST at room temperature for 2 h, and washed in PBST once, and then incubated with anti-PIWIL1 (Abcam, Cat# ab181056, 1:100 dilution), anti-hDcp1a (56-Y) antibodies (Santa Cruz Biotechnology, sc-100706, 1:500 dilution), and anti-UPF1 (Cell Signaling TECHNOLOGY, Cat# 12040, 1:200 dilution) in 3% BSA at 4 °C overnight. After incubation, cells were washed five times in PBST, 5 min each time. Cells were incubated with the secondary antibody FITC-conjugated AffiniPure goat anti-mouse IgG (H+L) (Jackson ImmunoResearch Laboratories, Cat# 115-095-003, 1:100 dilution) or FITC-conjugated AffiniPure goat anti-rabbit IgG (H+L) (Jackson ImmunoResearch Laboratories, Cat# 111-095-003, 1:100 dilution) or Alexa Fluor 594-conjugated AffiniPure goat anti-mouse IgG (H+L) (Jackson ImmunoResearch Laboratories, 115-585-003, Cat# 1:400 dilution) or Donkey anti-Rabbit IgG (H+L) Highly Cross-Adsorbed Secondary Antibody, Alexa Fluor 680 (ThermoFisher Scientific, Cat# A10043, 1:500 dilution) in PBST containing 3 % BSA at room temperature for 2 h, followed by a PBST wash once for 5 min. DAPI (Life Technologies, D1306, 1:5000 dilution) was then added to the PBST buffer and incubated at room temperature for 10 min, followed by three washes in PBST, 5 min each time. Coverslips were removed one at a time, and we added one drop of FluorPreserve™ (Merck/Millipore, Cat# 345787-25MLCN), mounted them to the glass slide, pressed gently, sealed them with nail polish, and stored them at 4°C overnight before confocal immunofluorescence microscopy (Zeiss, LSM710).

### PIWIL1 Knockout and knockdown Analyses

A pair of sgRNAs were cloned into a pGL3-U6-sgRNA-PGK-puromycin vector. pGL3-U6-sgRNA-PGK-puromycin and pST1374-N-NLS-flag-linker-Cas9-D10A were gifts from Xingxu Huang (Addgene plasmid # 51133; http://n2t.net/addgene:51133; RRID: Addgene: 51133). The knockout analysis was performed as previously described(86). sgRNAs and PIWIL1-knockout genome-PCR primers were listed in supplemental Table S5.

PIWIL1 Stealth siRNAs (Set of 3) HSS113746, HSS113747, HSS113748 were ordered from ThermoFisher Scientific. siRNAs were transfected into the cancer cells with Lipofectamine™ RNAiMAX Transfection Reagent (ThermoFisher Scientific, 13778150).

### Co-immunoprecipitation

Cells were lysed in co-immunoprecipitation lysis buffer [20mM Tris-HCl pH7.4, 150mM NaCl, 1 % (v/v) IGEPAL^®^ CA-630 (Millipore SIGMA, Cat# I8896), 1mM EDTA, 0.5 mM DTT, cOmplete™ EDTA-free Protease Inhibitor Cocktail Tablets (MERCK, Cat# 4693132001), and PhosStop Tablets (MERCK, 4906837001)], and spun at 14,000 rpm for 10 minutes to remove the debris. 10 μL of empty Dynabeads^®^ Protein G (ThermoFisher Scientific, Cat# 10004D) was added to the lysates and incubated for 0.5 hours for pre-clearing. 50 μL of empty Dynabeads^®^ Protein G was washed by Citrate-Phosphate Buffer (pH 5.0) and then incubated with 5ug Mouse Monoclonal anti-PIWIL1 antibody (Millipore SIGMA, Cat# SAB4200365) or 5ug Normal Mouse IgG Polyclonal Antibody (MERCK, Cat# 12-371). 50 μL of empty Dynabeads^®^ Protein A were washed by Citrate-Phosphate Buffer (pH 5.0) and then incubated with 15ul Rabbit Monoclonal anti-UPF1 antibody (Cell Signaling TECHNOLOGY, Cat# 12040) or 5ug Rabbit polyclonal anti-phospho-Upf1 (Ser1127) antibody (MERCK, Cat# 07-1016) or 5ug Rabbit polyclonal anti-UPF2 antibody (Abcam, ab157108) or 5ug Normal Rabbit IgG Polyclonal Antibody (MERCK, Cat# 12-370). All these incubations were done at 4°C for 4 hours with rotation. The Dynabeads Protein G/A-Ig complex was washed by IP wash buffer [20mM Tris-HCl pH7.4, 200mM NaCl, 0.05 % (v/v) IGEPAL^®^ CA-630, 0.5 mM DTT, cOmplete™ EDTA-free Protease Inhibitor Cocktail Tablets, and PhosStop Tablets] three times. The prewashed Dynabeads Protein G/A-Ig complex was incubated with the precleared lysates at 4°C overnight with rotation. The beads were then washed with IP wash buffer twice and IP high-salt wash buffer [20mM Tris-HCl pH7.4, 500mM NaCl, 0.05 % (v/v) IGEPAL^®^ CA-630, 0.5 mM DTT, cOmplete™ EDTA-free Protease Inhibitor Cocktail Tablets, and PhosStop Tablets] twice. Immunoprecipitates were run on 7.5% TGX Fast Cast acrylamide gels and probed with relevant antibodies for Western blotting or detected by using Coomassie Brilliant Blue staining solution based on the manufacturer’s protocol.

### Mass Spectrometry

The co-immunoprecipitated products were subjected to electrophoresis using 7.5% TGX Fast Cast acrylamide kit (Bio-Rad, 1610173TA) followed by Coomassie Brilliant Blue staining. Protein spots were then cut separately; the excised gel bands were first in-gel digested as previously describe(87). The obtained peptides were desalted, followed by nano-LC/MS/MS analysis.

Orbitrap Fusion MS equipped with nano EASY LC and Easyspray column (75 μm × 50 cm, PepMap RSLC C18 column, two μm, 100 Å, ThermoFisher Scientific) was employed to acquire MS data. The LC gradient of was 5 to 35% B (0.1% formic acid in CH3CN) in 60 min at a flow rate of 300 NL/min, the column temperate was set to 50 oC. The MS data were acquired in data-dependent mode. Briefly, a survey scan at 60K resolution is followed by ten HCD MS/MS scans performed in Orbitrap at 15K resolution.

### Ribonucleoprotein-immunoprecipitation (RIP)

Cells were homogenized in RIP lysis buffer [20mM Tris-HCl pH7.4, 150mM NaCl, 1 % (v/v) IGEPAL^®^ CA-630(Millipore SIGMA, Cat# I8896), 1mM EDTA, 0.5 mM DTT, cOmplete™ EDTA-free Protease Inhibitor Cocktail Tablets (MERCK, Cat# 4693132001), PhosStop Tablets (MERCK, 4906837001), and SUPERase• In™ RNase Inhibitor (ThermoFisher Scientific, Cat# AM2694)], and spun at 14,000 rpm for 10 minutes to remove the debris. 10 μL of empty Dynabeads^®^ Protein G (ThermoFisher Scientific, Cat# 10004D) was added to the lysates and incubated for 0.5 hours for pre-clearing. 50 μL of empty Dynabeads^®^ Protein G was washed by Citrate-Phosphate Buffer (pH 5.0) and then incubated with 5ug Mouse Monoclonal anti-PIWIL1 antibody (Millipore SIGMA, Cat# SAB4200365) for 4 hours at 4°C with rotation. 50 μL of empty Dynabeads^®^ Protein A was washed by Citrate-Phosphate Buffer (pH 5.0) and then incubated with 15ul Rabbit Monoclonal anti-UPF1 antibody (Cell Signaling Technology, Cat# 12040) or 5ug Normal Rabbit IgG Polyclonal Antibody (Merck, Cat# 12-370). All these incubations were done at 4°C for 4 hours with rotation. The Dynabeads Protein G/A-Ig complex was washed by RIP wash buffer [20mM Tris-HCl pH7.4, 200mM NaCl, 0.05 % (v/v) IGEPAL^®^ CA-630, 0.5 mM DTT, and SUPERase• In™ RNase Inhibitor] three times. The prewashed Dynabeads Protein G/A-Ig complex was incubated with the precleared lysates at 4°C overnight with rotation. The beads were then washed with RIP wash buffer twice and RIP high-salt wash buffer [20mM Tris-HCl pH7.4, 500mM NaCl, 0.05 % (v/v) IGEPAL^®^ CA-630, 0.5 mM DTT, and SUPERase• In™ RNase Inhibitor] twice and then the RNA was eluted using 1 mL of TRIzol^®^ (ThermoFisher Scientific, Cat# 15596026) following the manufacturer’s manual.

### Small RNA-Seq library construction and sequencing

Small RNA libraries were prepared from the extracted 1ug total RNA or 100ng PIWIL1-antibody-pulled down RNA using NEBNext Multiplex Small RNA Library Prep Set for Illumina (Set 1) (NEB, Cat# E7300S) according to the manufacture’s protocol. The libraries were sequenced using the Illumina HiSeqX platform for 150-nucleotide pair-end runs.

### β-elimination by NaIO_4_-Oxidation and Small RNA-Seq library construction

20ug total RNA was oxidized to achieve β-elimination by exposing to 20 mM NaIO_4_ (Sigma-Aldrich, Cat# 311448) in 200 mM lysine-HCl buffer (pH 8.5, Sigma-Aldrich, Cat# 62929) in a total volume of 40 μl at 37 °C for 30 min with shaking in a dark tube. The reaction was quenched by 2 μl ethylene glycol. The RNA was column-purified (RNA Clean & Concentrator-5, Zymo, Cat# R1015) and eluted in 8 μl molecular biology grade, RNase-free water. 6ul eluted RNA was used for Small RNA libraries using NEBNext Multiplex Small RNA Library Prep Set for Illumina (Set 1) (NEB, Cat# E7300S) according to the manufacture’s protocol. The libraries were sequenced using the Illumina HiSeqX platform for 150-nucleotide pair-end runs.

### Analyses of small RNA sequencing data

Small RNA sequences were processed with TrimGalore (version 0.4.4_dev; http://www.bioinformatics.babraham.ac.uk/projects/trim_galore/) with the default adapter trimming mode to auto-detect adapters in the 150bp long sequencing reads. A size cutoff using 17bp to 42 bp was applied to retain small RNA reads of suitable length. The commands were: “trim_galore --phred33 --gzip -q 20 --fastqc --output_dir trimmed_fqc_out --length 17 --max_length 42 --trim-n $read”. Subsequently trimmed reads were mapped onto the corresponding genome [human (hg38) or mouse (mm10)] using bowtie1 (version 1.2.1.1)(88)with commands “bowtie -S -p 5 -v 0 -n 0 -l 18 -k 1 --no-unal --al $GMAPPED_FQ *Homo_sapiens*. GRCh38.dna.primary_assembly.fa $read > $GMAPPED_SAM”. Genome-mapped reads were sequentially mapped to the libraries of miRNA, tRNA, rRNA, snRNA, snoRNA, piRNAs, and repetitive elements. miRNA libraries were obtained from miRBase (March 11, 2018)(89). tRNA reference fasta was built by combining the tRNA sequences from GtRNAdb(90) and those from genome annotation. rRNA libraries were prepared from RNAcentral (January of 2019)(91). Both snRNA and snoRNA libraries were obtained from both genome annotation and RNAcentral (January of 2019). piRNAs were classified as *de novo* and known, with the former identified by proTRAC (version 2.4.2)(92), and the latter by mapping onto piRNAs from piRNABank. Commands for proTRAC were: “perl proTRAC_2.4.2.pl -genome Homo_sapiens. GRCh38.dna.primary_assembly.header.simple.fa -map $map_file -repeatmasker hg38.repeatMasker.matched.gtf -geneset Homo_sapiens.GRCm38.95.chr.gtf -pimin 25 -pimax 31”. $mapfile was the mapped output using sRNAmapper (version 1.0.4) (93). Repeat-derived small RNA libraries were obtained from the RepeatMasker annotation in UCSC genome browser.

### RIP-Seq data analysis

Each of the strand-specific RNA-Seq samples was first subject to a quality trimming step using TrimGalore (0.4.4_dev) to remove low quality bases (retaining only Q20 nucleotides) and reads shorter than 50 nucleotides using TrimGalore (0.4.4_dev) (trim_galore --fastqc --length 50 --trim-n --paired $R1 $R2). Adapter sequences were absent as the read counts were long as 150 bp, but were still evaluated using FastQC (v0.11.5) to ensure the absence of adapters (fastqc -o $out_dir -f fastq -t 10 $R1.fastq.gz $R2.fastq.gz). Then the paired-end reads were mapped onto the human genome (hg38) with STAR (version 020201) (94) to obtain mapped BAM files (STAR --runMode alignReads --runThreadN 8 --genomeDir $hg38_dir --sjdbGTFfile Homo_sapiens.GRCh38.89.gtf --readFilesCommand zcat --readFilesIn $trimmed.R1.fq.gz $trimmed.R2.fq.gz --outSAMtype BAM SortedByCoordinate --outFileNamePrefix $sample). Subsequently, htseq-count (version 0.9.0) with strand-bias option (htseq-count -f bam -s reverse sample.sortedByCoord.out.bam (58, 62) Homo_sapiens.GRCh38.89.gtf > sample.htseq.counts) was used to obtain a read count estimate for each annotated gene of the human genome. The raw read counts for each sample were used for following differential expression analyses to identify mRNAs bound by PIWIL1 and UPF1.

To identify PIWIL1-bound genes, we used an RNA immunoprecipitation followed by deep sequencing (RIP-Seq). The PIWIL1-antibody was used to pull down putative PIWIL1-bound mRNAs in SNU-1 cell lines, which were subject to a library preparation protocol for strand-specific transcriptome sequencing using RNA-Seq. This sample was denoted as “WT_PIWIL1-RIP.” To measure the binding strength between PIWIL1 protein and its bound mRNAs, an input sample without using an antibody for immunoprecipitation (namely the mRNAs from SNU-1 cell line) was prepared and defined as “WT-input.” To remove possible false positives, RNA immunoprecipitation and input were also prepared for the PIWIL1 knock-out cell lines of SNU-1, for which the samples were denoted as “KO_PIWIL1_RIP” and “KO-input.” Two biological replicates were generated for each of the four samples. To identify PIWIL1-bound mRNAs, differential expression analysis was performed between the input-normalized WT PIWIL1-RIP raw read counts and the input-normalized PIWIL-KO-RIP raw counts using the DESeq2 package (95) in R. In particular, firstly, for the eight samples, the size factor (level 1, involving eight values across the eight samples) function in DESeq2 package in R was applied to obtain the normalized counts in each sample. Subsequently, keeping the counts for PIWIL1-KO-RIP samples unchanged, a size factor between the WT PIWIL1 input and PIWIL1-KO input (level 2, involving 2 values across two replicates of “input” samples) was calculated for each of the two replicates, respectively. This size factor (for each of the two replicates) was then used to normalize the counts for the two WT PIWIL1 RIP replicates, respectively. After this normalization with the input read counts, the WT PIWIL1 RIP, and PIWIL1-KO RIP samples were directly comparable as the influence from the input was already removed. Then, the normalized read counts for each of the RIP samples (two WT PIWIL1 RIP and two PIWIL1-KO RIP) were converted back to raw read counts using the initial size factors (level 1). These raw read counts between the WT PIWIL1 RIP and PIWIL1-KO RIP, which have been already normalized by input for each respective sample, were used for DE analyses to identify DE features between the WT_RIP and KO_RIP samples. A P value cutoff of 0.05 after applying an adjustment of multiple comparisons and a fold change cutoff of 1.5 were considered as PIWIL1-bound genes.

To identify UPF1-bound genes in the PIWIL1-positive background, we directly compared the significantly upregulated RNAs in UPF1 immunoprecipitated mRNAs (WT UPF1 RIP) versus IgG immunoprecipitated mRNAs (WT IgG RIP) in SNU-1 cell line (each containing two biological replicates) was used. Differential expression analysis was directly performed between UPF1 RIP and IgG RIP samples with DEseq2 package in R. Similarly, an adjusted P-value cutoff of 0.05 and a fold change cutoff of 1.5 was used to identify UPF1-bound features in the SNU-1 cell line.

### Signed weighted gene co-expression network analysis (WGCNA)

We produced five independent batches of RNA-Seq data with 10 samples from two groups. WT-group: WT1 (WT 1^st^ batch), WT2 (KO-Con 2^nd^ batch), WT3 (WT 3^rd^ batch), WT4 (KO-Con 4^th^ batch), WT5 (WT 5^th^ batch); KO group: KO1 (KO-#20 1^st^ batch), KO2 (KO-#37, 2^nd^ batch), KO3 (KO-#20 3^rd^ batch), KO4 (KO-#37 4^th^ batch), KO5 (KO-#20, 5^th^ batch)) for RNA-Seq WGCNA data analysis with NovelBrain Cloud Analysis Platform. Briefly, after the libraries were quality controlled with a Bioanalyzer 2200 (Agilent, Santa Clara, CA) and sequenced by HiSeq X (Illumina, San Diego, CA) on a 150 bp paired-end run, clean reads were obtained from the raw reads by removing the adaptor sequences, and low-quality reads. Then, the clean reads were then aligned to the human genome (GRCh38, NCBI) using the Hisat2 (96). Htseq (97) was used to get gene counts, and the RPKM method was used to determine the gene expression. WGCNA (98) was performed across all samples using the standard method with a power of 17 to cluster the expression patterns. WGCNA was used to construct a network in published data sets independently and generated an independent list of hub genes (kME > 0.9) for each data set. RUVseq algorithm was used for batch-to-batch correction.

## ACKNOLEDGEMENTS

We are grateful to Dr. Peixiang Ma for doing the PIWIL1-UPF1 docking analysis, Dr. Jiangsha Zhao, Ms. Zhaoran Zhang, Weidong Chen, and Jiawei Zhang for various technical assistance, Prof. Bin Shen and Prof. Xingxu Huang for offering us CRISPR-Cas9 KO system. We thank Yuedong Huang, Kun-Yong Kim, Nils Neuenkirchen, Yiying Yang, Chen Wang, Zukai Liu, Tingting Lu, and Yuanyuan Gong for comments on the manuscript. This work was supported by ShanghaiTech University.

