## Supplementary Figures and Tables for "PIWIL1 Promotes Gastric Cancer via a piRNA-Independent Mechanism"

SI Appendix

Table of Contents

| | |
|---|---|
| Supplementary Figure S1-6 Legends | pp 1-6 |
| Supplementary Figs. S1-6 | pp 7-12 |
| Supplementary Tables S1-5 | pp 13-17 |

### Supplementary Figure Legends

**Fig. S1.** The expression of PIWIL2 and PIWIL4 in gastric clinical samples and gastric cancer cell lines.

(A) Bar graph showing quantitative RT-PCR of PIWIL2 and PIWIL4 mRNA expression in one normal gastric epithelial cell line GES-1 and six human gastric cancer cell lines. The mRNA level of GES-1 was normalized to 1. β-Actin was used as an internal standard to normalize the level of PIWIL2 or PIWIL4 from the same sample.

(B) Western blot showing protein levels of PIWIL1 in GES-1 and six human gastric cancer cell lines. β-Actin was used as a loading control.

(C) Analysis of mRNA expression of all human PIWI family members in human gastric cancer tissues and normal gastric tissues by using the Cho gastric dataset of the ONCOMINE database.

(D) Analysis of PIWIL1 mRNA expression in human gastric cancer using the Cho gastric dataset of the ONCOMING database. Data are grouped by cancer types.

(E) Analysis of correlation between PIWIL1 mRNA expression and the survival of different types of gastric cancer patients. The data source is the KM-plotter database. Patients are split by the automatic select best cutoff. P: log-rank p; HR: hazard ratio.

**Fig. S2** Knocking-down PIWIL1 significantly inhibits cell cycle and migration in AGS cells.

(A) Diagram of CRISPR-Cas9 nickase knockout system with one pair of sgRNAs of PIWIL1.

(B) Western blot analysis of the PIWIL1 protein expression in PIWIL1-WT and PIWIL1-KO SNU-1 cells. WT: wildtype SNU-1 cells without transfection of CRISPR-Cas9 plasmids; KO-Con: SNU-1 cells co-transfected with PIWIL1-sgRNA plasmid and Cas9 plasmid but gene-editing did not occur in *PIWIL1*; KO-#20, KO-#37, and KO-#66: three independent SNU-1 cell clones which were co-transfected with PIWIL1-sgRNA plasmid and Cas9 plasmid with gene-editing occurred in *PIWIL1* to knockout PIWIL1 protein.

(C) Apoptosis assay of PIWIL1-WT or PIWIL1-KO SNU-1 cells using Annexin V-FITC and 7-amino-Actinomycin-D (7-AAD) to label early and late apoptotic cells, respectively.

(D) Bar graph showing numbers of the apoptotic cell of PIWIL1-WT and PIWIL1-KO SNU-1 cells.

(E) The PIWIL1 protein expression in AGS cells treated with PIWIL1-siRNA-Control (si-control), PIWIL1-siRNA-#46 (si-PIWIL1-#46), PIWIL1-siRNA-#47 (si-PIWIL1-#47) or PIWIL1-siRNA-#48 (si-PIWIL1-#48).

(F) Transwell assay of si-control, si-PIWIL1-#46, si-PIWIL1-#47, or si-PIWIL1-#48 AGS cells.

(G) Bar graph showing the numbers of migrated cells in the transwell assay.

(H) Flow cytometry analysis of cell cycle of si-control, si-PIWIL1-#46, si-PIWIL1-#47, or si-PIWIL1-#48 AGS cells.

(I) Bar graph showing the percentage of the G1 and S phase of si-control, si-PIWIL1-#46, si-PIWIL1-#47, or si-PIWIL1-#48 AGS cells.

**Fig. S3** Changes of RNA co-expression modules and PIWIL1-target RNAs in PIWIL1-KO SNU-1 cells.

(A) Heatmap of the correlation between module eigengenes and the trait of with or without PIWIL1 expression, continued from Fig 3B, showing 10 additional modules with correlation to *PIWIL1-KO*. Red color represents a positive correlation between a module and the trait, and blue color represents a negative correlation. Each cell contained the corresponding correlation and *P-value*. Note that the green module shows a significant negative correlation ($P$=0.00014). The remaining 11 modules show no correlation and are not included in this paper.

(B) Bar plot of mean gene significance across gene co-expression modules from WGCNA analysis of wildtype (WT) and PIWIL1-KO SNU-1 cells.

(C) The histograms displaying the eigengene expression of the blue or turquoise modules from ten WT or PIWIL1-KO samples, which indicated good repeatability eigengene (ME) between the samples in either WT or KO group within each modules.

(D) Quantitative RT-PCR validation of the expression of 8 well-studied oncogenic mRNA from the blue module and 10 tumor suppressor genes from turquoise modules.

(E) Western blotting of PIWIL1-RIP of WT and PIWIL1-KO SNU-1 cells using 10% of the immunoprecipitate to confirm the antibody IP efficiency. The remaining 90% were used for RNA extraction and deep sequencing.

(F) Distribution of different types of PIWIL1-bound RNAs.

(G) Scatter chart illustrating the differently expressed protein-coding (left) or long-noncoding (middle) or pseudogene (right) RNAs in the PIWIL1-KO cells (blue dots). Red circles are the PIWIL1-bound RNAs that are differentially expressed in PIWIL1-KO cells.

(H) KEGG pathway analysis of the 18 PIWIL1-bound mRNAs that are positively regulated by PIWIL1, showing enrichment in the glycolysis/gluconeogenesis pathway.

(I) Venn diagram illustrating the overlap between RNAs that are not bound by PIWIL1 but negatively regulated PIWIL1 that are not bound by PIWIL1 but positively regulated PIWIL1, and the blue-module hub genes, with kME≥0.9, P<0.05 and fold-change≥1.5

cutoff.

(J) KEGG pathway analysis of the 221 PIWIL1 positively and indirectly regulated hub genes. The cell cycle pathway is significantly enriched in these genes.

**Fig. S4** There are abundant piRNAs bound by PIWIL1 in mouse testis.

(A) Size profiles of each class of small RNAs immunoprecipitated by anti-PIWIL1 antibody in mouse testes, with IgG as the negative IP control.

(B) The expression (indicated as Transcripts Per Million, TPM) of key components of the germline piRNA pathway by RNA-Seq in WT SNU-1 cells; genes were grouped according to their roles in the piRNA pathway. Gene expression data were calculated from triplicates.

**Fig. S5** PIWIL1 interacts with the NMD complex in P body in gastric cancer cell line AGS.

(A) Western blotting showing that NMD complex core proteins UPF1, UPF2, and phosphorylated UPF1 can be pulled down by the PIWIL1 antibody in AGS gastric cancer cells.

(B) Immunofluorescence staining of PIWIL1 (green) and DCP1A (red, P body marker) in AGS cells.

(C) The 203 PIWIL1-negatively regulated RNAs, which are targeted by both PIWIL1 and UPF1, were analyzed by the Gene Ontology (GO) analysis of biological process.

**Fig. S6** The expression of UPF1 in gastric cancer clinical samples and the correlation between PIWIL1 and UPF1 mRNA expressions.

(A) UPF1 mRNA levels in 97 gastric cancer patients examined by quantitative RT-PCR, using β-Actin as an internal control. Bar value ($\log_2$) represents the difference of UPF1 mRNA levels between normal tissue and tumor. Bar-value > 1 indicates that UPF1 mRNA levels increase ≥2-fold in tumors. Bar-value <-1 indicates that UPF1 mRNA levels are ≥2-fold lower in tumors. Data were calculated from triplicates.

(B) GEPIA database (http://gepia.cancer-pku.cn/) shows that UPF1 mRNA expression in gastric cancer samples is higher than that in normal tissue samples. There are 407 gastric cancer samples and 211 normal tissue samples in this chart.

(C) Scatter plot showing the correlation between PIWIL1 and UPF1 mRNA expressions in 97 pairs of gastric cancer and paired normal samples. Pearson test was used to evaluate the significance of correlation.

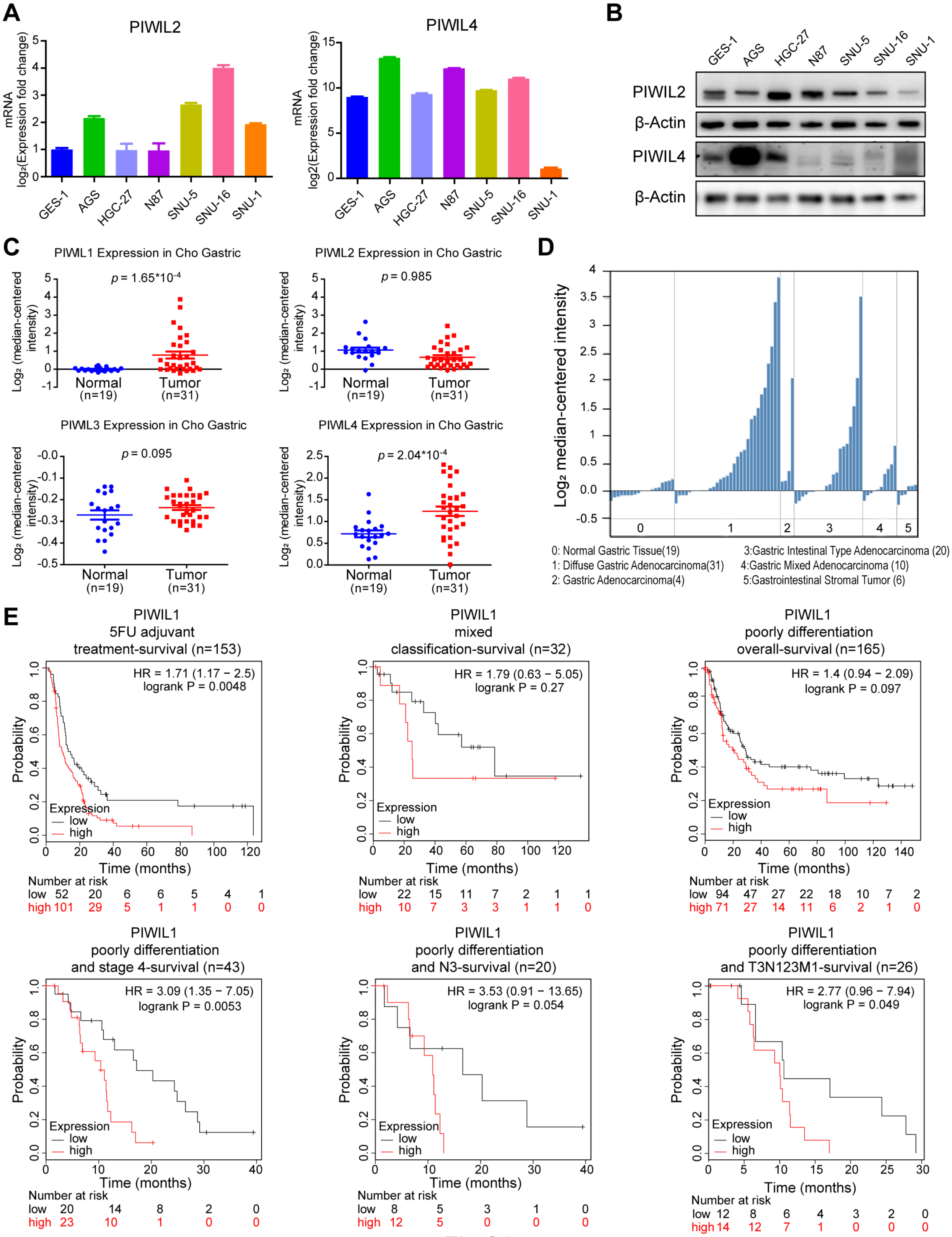
A
PIWIL2
PIWIL4
mRNA log₂(Expression fold change)
GES-1
AGS
HGC-27
N87
SNU-5
SNU-16
SNU-1
B
PIWIL2
β-Actin
PIWIL4
β-Actin
C
PIWL1 Expression in Cho Gastric
PIWL2 Expression in Cho Gastric
PIWL3 Expression in Cho Gastric
PIWL4 Expression in Cho Gastric
Log₂ (median-centered intensity)
p = 1.65*10-4
p = 0.985
p = 0.095
p = 2.04*10-4
Normal (n=19)
Tumor (n=31)
D
Log₂ median-centered intensity
0: Normal Gastric Tissue(19)
1: Diffuse Gastric Adenocarcinoma(31)
2: Gastric Adenocarcinoma(4)
3:Gastric Intestinal Type Adenocarcinoma (20)
4:Gastric Mixed Adenocarcinoma (10)
5:Gastrointestinal Stromal Tumor (6)
E
PIWIL1 5FU adjuvant treatment-survival (n=153)
HR = 1.71 (1.17 − 2.5) logrank P = 0.0048
PIWIL1 mixed classification-survival (n=32)
HR = 1.79 (0.63 − 5.05) logrank P = 0.27
PIWIL1 poorly differentiation overall-survival (n=165)
HR = 1.4 (0.94 − 2.09) logrank P = 0.097
PIWIL1 poorly differentiation and stage 4-survival (n=43)
HR = 3.09 (1.35 − 7.05) logrank P = 0.0053
PIWIL1 poorly differentiation and N3-survival (n=20)
HR = 3.53 (0.91 − 13.65) logrank P = 0.054
PIWIL1 poorly differentiation and T3N123M1-survival (n=26)
HR = 2.77 (0.96 − 7.94) logrank P = 0.049
Probability
Time (months)
Expression
low
high
Number at risk
low 52 20 6 6 5 4 1
high 101 29 5 1 1 0 0
low 22 15 11 7 2 1 1
high 10 7 3 3 1 1 0
low 94 47 27 22 18 10 7 2
high 71 27 14 11 6 2 1 0
low 20 14 8 2 0
high 23 10 1 0 0
low 8 5 3 1 0
high 12 5 0 0 0
low 12 8 6 4 3 2 0
high 14 12 7 1 0 0 0

Fig.S1

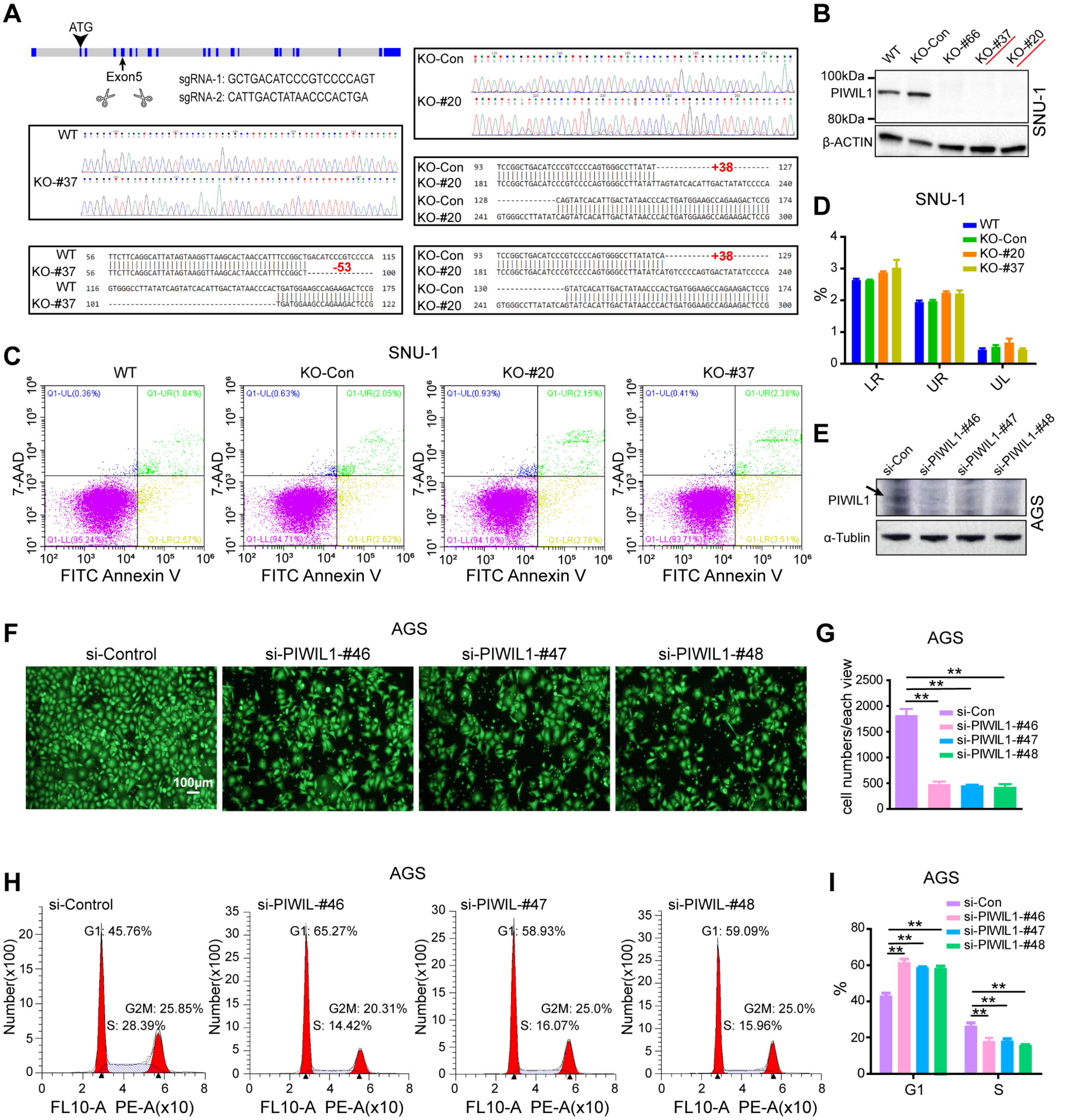
A
ATG
Exon5
sgRNA-1: GCTGACATCCCGTCCCCAGT
sgRNA-2: CATTGACTATAACCCACTGA
WT
KO-#37
KO-Con
KO-#20
+38
-53
B
WT
KO-Con
KO-#66
KO-#37
KO-#20
100kDa
PIWIL1
80kDa
β-ACTIN
SNU-1
C
SNU-1
WT
KO-Con
KO-#20
KO-#37
7-AAD
FITC Annexin V
D
SNU-1
WT
KO-Con
KO-#20
KO-#37
%
LR
UR
UL
E
si-Con
si-PIWIL1-#46
si-PIWIL1-#47
si-PIWIL1-#48
PIWIL1
α-Tublin
AGS
F
AGS
si-Control
si-PIWIL1-#46
si-PIWIL1-#47
si-PIWIL1-#48
100μm
G
AGS
cell numbers/each view
si-Con
si-PIWIL1-#46
si-PIWIL1-#47
si-PIWIL1-#48
H
AGS
si-Control
G1: 45.76%
G2M: 25.85%
S: 28.39%
si-PIWIL-#46
G1: 65.27%
G2M: 20.31%
S: 14.42%
si-PIWIL-#47
G1: 58.93%
G2M: 25.0%
S: 16.07%
si-PIWIL-#48
G1: 59.09%
G2M: 25.0%
S: 15.96%
Number(x100)
FL10-A PE-A(x10)
I
AGS
si-Con
si-PIWIL1-#46
si-PIWIL1-#47
si-PIWIL1-#48
%
G1
S

Fig.S2

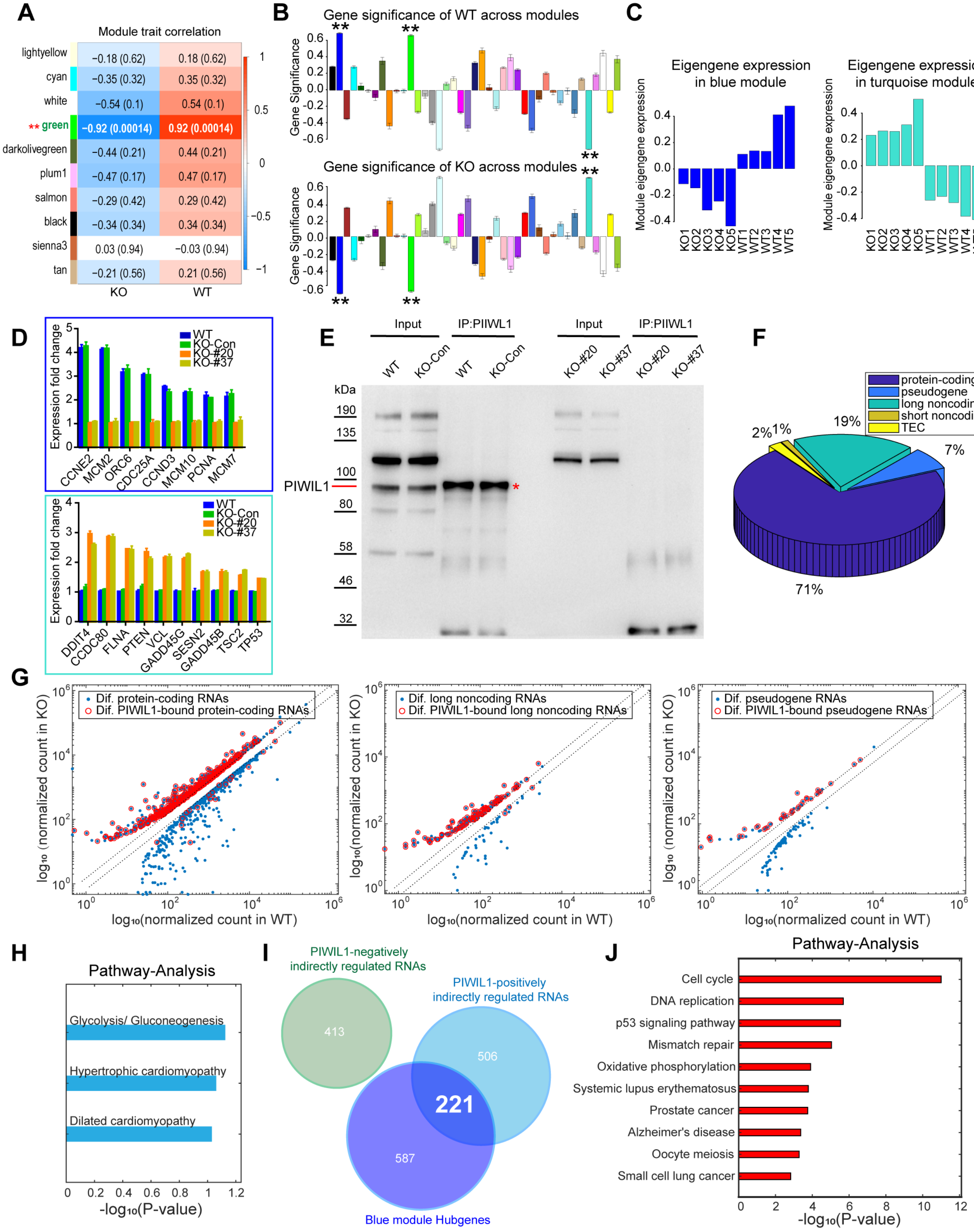

A
Module trait correlation
lightyellow
cyan
white
** green
darkolivegreen
plum1
salmon
black
sienna3
tan
−0.18 (0.62)
0.18 (0.62)
−0.35 (0.32)
0.35 (0.32)
−0.54 (0.1)
0.54 (0.1)
−0.92 (0.00014)
0.92 (0.00014)
−0.44 (0.21)
0.44 (0.21)
−0.47 (0.17)
0.47 (0.17)
−0.29 (0.42)
0.29 (0.42)
−0.34 (0.34)
0.34 (0.34)
0.03 (0.94)
−0.03 (0.94)
−0.21 (0.56)
0.21 (0.56)
KO
WT
B
Gene significance of WT across modules
Gene significance of KO across modules
Gene Significance
C
Eigengene expression in blue module
Eigengene expression in turquoise module
Module eigengene expression
KO1 KO2 KO3 KO4 KO5 WT1 WT2 WT3 WT4 WT5
D
Expression fold change
WT
KO-Con
KO-#20
KO-#37
CCNE2 MCM2 ORC6 CDC25A CCND3 MCM10 PCNA MCM7
DDIT4 CCDC80 FLNA PTEN VCL GADD45G SESN2 GADD45B TSC2 TP53
E
Input
IP:PIWIL1
WT
KO-Con
KO-#20
KO-#37
kDa
190
135
100
80
58
46
32
PIWIL1
F
protein-coding
pseudogene
long noncoding
short noncoding
TEC
2%
1%
19%
7%
71%
G
Dif. protein-coding RNAs
Dif. PIWIL1-bound protein-coding RNAs
Dif. long noncoding RNAs
Dif. PIWIL1-bound long noncoding RNAs
Dif. pseudogene RNAs
Dif. PIWIL1-bound pseudogene RNAs
log10 (normalized count in KO)
log10(normalized count in WT)
H
Pathway-Analysis
Glycolysis/ Gluconeogenesis
Hypertrophic cardiomyopathy
Dilated cardiomyopathy
-log10(P-value)
I
PIWIL1-negatively indirectly regulated RNAs
PIWIL1-positively indirectly regulated RNAs
Blue module Hubgenes
413
506
221
587
J
Pathway-Analysis
Cell cycle
DNA replication
p53 signaling pathway
Mismatch repair
Oxidative phosphorylation
Systemic lupus erythematosus
Prostate cancer
Alzheimer's disease
Oocyte meiosis
Small cell lung cancer
-log10(P-value)

Fig.S3

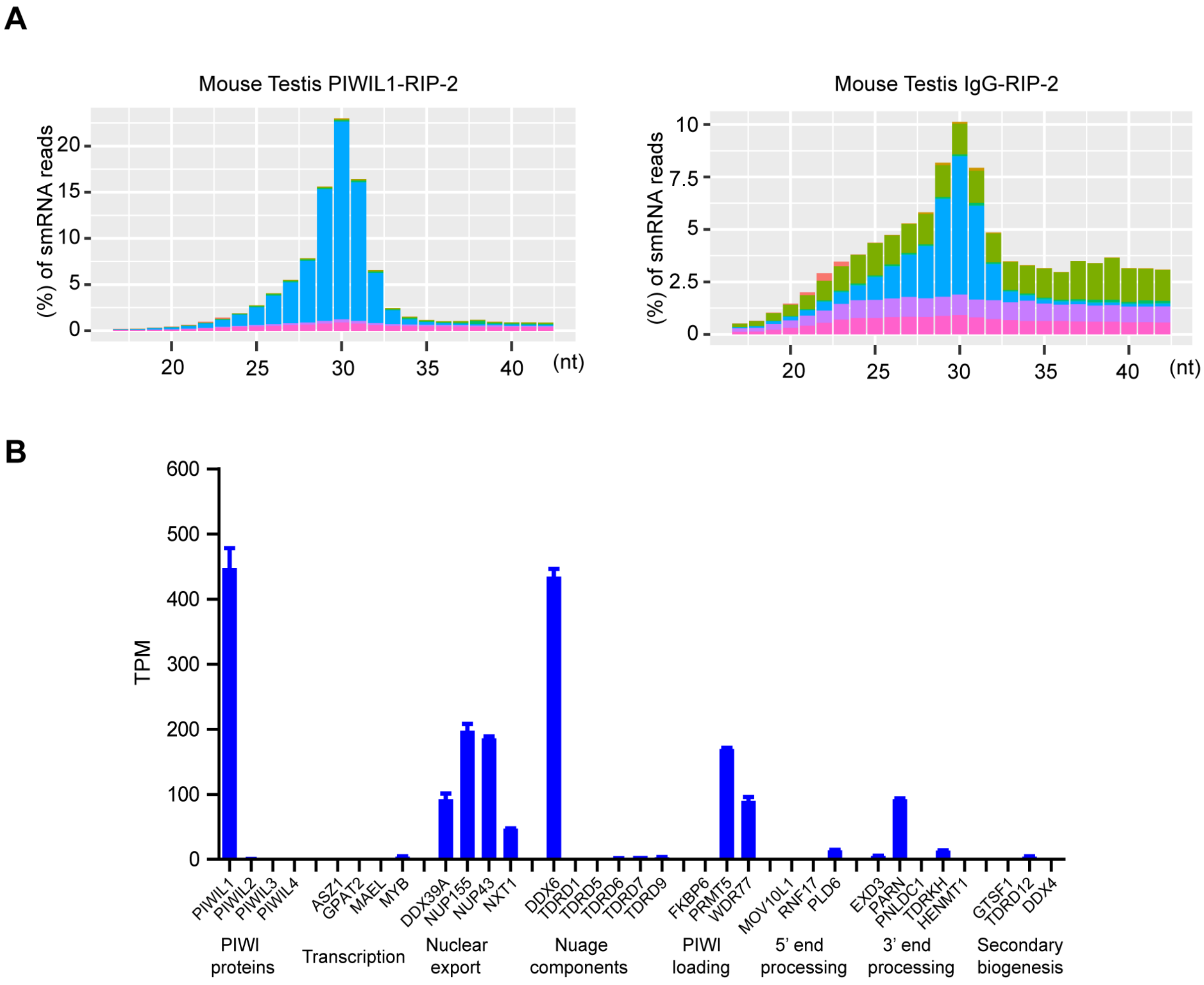

A
Mouse Testis PIWIL1-RIP-2
(%) of smRNA reads
0
5
10
15
20
20
25
30
35
40
(nt)
Mouse Testis IgG-RIP-2
(%) of smRNA reads
0
2.5
5
7.5
10
20
25
30
35
40
(nt)
B
TPM
0
100
200
300
400
500
600
PIWIL1
PIWIL2
PIWIL3
PIWIL4
ASZ1
GPAT2
MAEL
MYB
DDX39A
NUP155
NUP43
NXT1
DDX6
TDRD1
TDRD5
TDRD6
TDRD7
TDRD9
FKBP6
PRMT5
WDR77
MOV10L1
RNF17
PLD6
EXD3
PARN
PNLDC1
TDRKH
HENMT1
GTSF1
TDRD12
DDX4
PIWI proteins
Transcription
Nuclear export
Nuage components
PIWI loading
5' end processing
3' end processing
Secondary biogenesis

Fig.S4

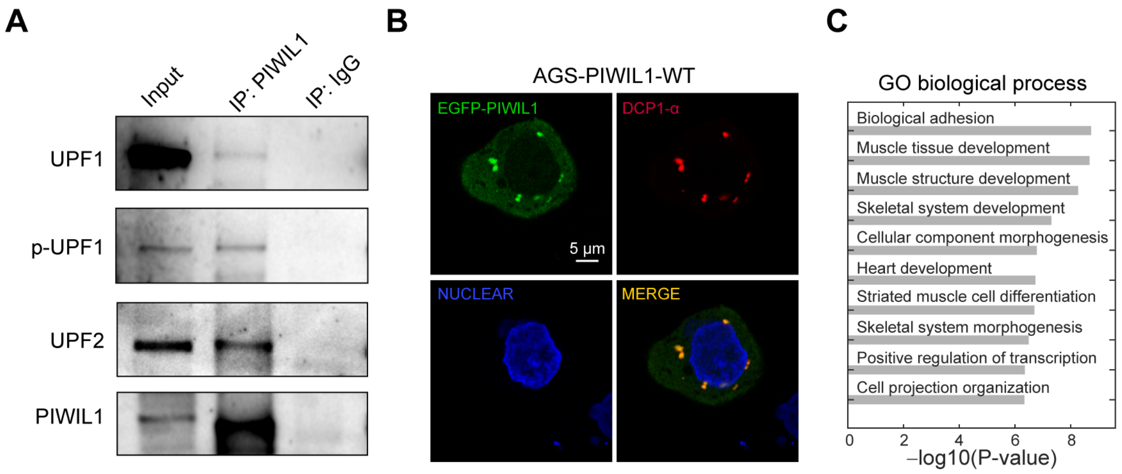
A
Input
IP: PIWIL1
IP: IgG
UPF1
p-UPF1
UPF2
PIWIL1
B
AGS-PIWIL1-WT
EGFP-PIWIL1
DCP1-α
5 μm
NUCLEAR
MERGE
C
GO biological process
Biological adhesion
Muscle tissue development
Muscle structure development
Skeletal system development
Cellular component morphogenesis
Heart development
Striated muscle cell differentiation
Skeletal system morphogenesis
Positive regulation of transcription
Cell projection organization
0
2
4
6
8
–log10(P-value)

Fig.S5

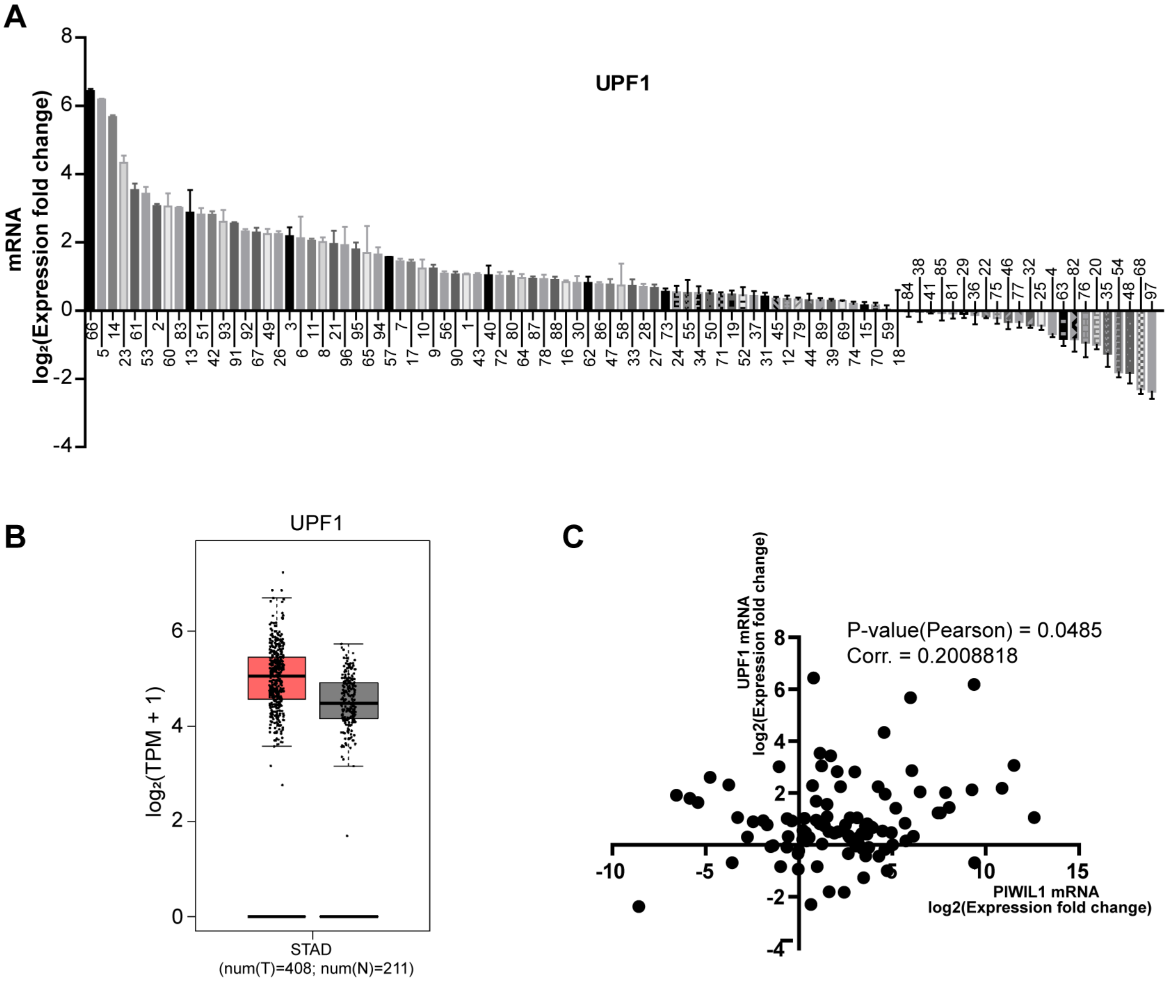
A
UPF1
mRNA
log₂(Expression fold change)
B
UPF1
log₂(TPM + 1)
STAD
(num(T)=408; num(N)=211)
C
P-value(Pearson) = 0.0485
Corr. = 0.2008818
UPF1 mRNA
log2(Expression fold change)
PIWIL1 mRNA
log2(Expression fold change)

Fig.S6

**Table S1** Relationship between levels of PIWIL1 expression in gastric cancer clinical samples and pathological features.

Grade of tumor differentiation and metastasis was determined by the pathologists. Tumor stages were classified according to the tumor node metastasis (TNM) classification of the American Joint Committee on Cancer and the International Union. PIWIL1 positive: PIWIL1 mRNA expression of tumor tissue is higher than that of paired normal tissue. PIWIL1 negative: PIWIL1 mRNA expression of tumor tissue is equal or lower than that of paired normal tissue.

| clinical characteristics | No.patients | PIWIL1 postive(n=74) | PIWIL1 negative(n=23) | Spearman's correlation |
|---|---|---|---|---|
| **Gender** | | | | |
| Male | 68 | 54 | 14 | p=0.1333 |
| Female | 29 | 20 | 9 | rho=0.1536 |
| **Metastasis** | | | | |
| N0 | 26 | 15 | 11 | p=0.00189 |
| N1 | 19 | 16 | 3 | rho=0.3116101 |
| N2 | 14 | 13 | 1 | |
| N3 | 31 | 23 | 8 | |
| M1 | 7 | 7 | 0 | |
| **Tumor differentiation** | | | | |
| Poor | 54 | 46 | 8 | p=0.002889 |
| Moderate | 38 | 28 | 10 | rho=-0.2994173 |
| High | 5 | 0 | 5 | |
| **TNM stage** | | | | |
| I | 5 | 0 | 5 | p=0.0007306 |
| II | 25 | 16 | 9 | rho=0.3372339 |
| III | 56 | 47 | 9 | |
| IV | 11 | 11 | 0 | |

**Table S2** Mapped reads of each class of small RNAs in individual libraries in the mouse testis and human gastric cancer cell line SNU-1.

| Sample | miRNA | tRNA | rRNA | snRNA | snoRNA | piRNA | Repeat | other_smRNA |
|---|---|---|---|---|---|---|---|---|
| Mouse Testis_total_smRNA | 12678507 | 12678507 | 12678507 | 12678507 | 12678507 | 12678507 | 12678507 | 12678507 |
| Mouse Testis_NaIO4-Oxidation | 14900569 | 14900569 | 14900569 | 14900569 | 14900569 | 14900569 | 14900569 | 14900569 |
| Mouse Testis IgG-RIP-1 | 79018 | 72239 | 3336412 | 173077 | 93570 | 2939996 | 1935068 | 1648942 |
| Mouse Testis IgG-RIP-2 | 59090 | 54597 | 2540694 | 131836 | 70570 | 2218710 | 1478321 | 1253337 |
| Mouse Testis PIWIL1-RIP-1 | 56730 | 87998 | 544530 | 605729 | 38018 | 18499860 | 933020 | 2674728 |
| Mouse Testis PIWIL1-RIP-2 | 40396 | 60987 | 377844 | 416802 | 26318 | 12762165 | 648836 | 1858764 |
| WT_total_smRNA | 12296858 | 12296858 | 12296858 | 12296858 | 12296858 | 12296858 | 12296858 | 12296858 |
| KO-control_total_smRNA | 12963083 | 12963083 | 12963083 | 12963083 | 12963083 | 12963083 | 12963083 | 12963083 |
| WT_PIWIL1-RIP | 139062 | 229791 | 8030386 | 713548 | 113122 | 37091 | 662167 | 2795939 |
| KO-control_PIWIL1-RIP | 153312 | 251663 | 8756673 | 783108 | 124193 | 36167 | 726704 | 3066667 |
| WT_NaIO4-Oxidation | 3972527 | 7347060.551 | 6181521.806 | 2908260 | 29662406 | 599283 | 2338048 | 4594911 |
| KO-#20_total_smRNA | 17187053 | 17187053 | 17187053 | 17187053 | 17187053 | 17187053 | 17187053 | 17187053 |
| KO-#37_total_smRNA | 11071450 | 11071450 | 11071450 | 11071450 | 11071450 | 11071450 | 11071450 | 11071450 |
| KO-#20_PIWIL1-RIP | 70139 | 144583 | 9898712 | 596265 | 228979 | 40393 | 617673 | 2521023 |
| KO-#37_PIWIL1-RIP | 77562 | 165396 | 11314274 | 681244 | 263496 | 59578 | 703619 | 2859300 |
| KO-#20_NaIO4-Oxidation | 4523005 | 8111631.373 | 7606573.73 | 1337391 | 12489727 | 521420 | 2718902 | 4153131 |

**Table S3** The proportion of each class of small RNA libraries in the mouse testis and human gastric cancer cell line SNU-1.

| Sample | miRNA | tRNA | rRNA | snRNA | snoRNA | piRNA | Repeat | Other_smRNA |
|---|---|---|---|---|---|---|---|---|
| Mouse Testis_total_smRNA | 7.496% | 0.587% | 2.804% | 0.884% | 0.550% | 79.646% | 2.378% | 5.655% |
| Mouse Testis_NaIO4-Oxidation | 0.553% | 0.538% | 1.545% | 1.029% | 0.596% | 91.198% | 1.411% | 3.130% |
| Mouse Testis IgG-RIP-1 | 0.769% | 0.703% | 32.461% | 1.684% | 0.910% | 28.604% | 18.827% | 16.043% |
| Mouse Testis IgG-RIP-2 | 0.757% | 0.699% | 32.543% | 1.689% | 0.904% | 28.419% | 18.935% | 16.054% |
| Mouse Testis PIWIL1-RIP-1 | 0.242% | 0.375% | 2.323% | 2.584% | 0.162% | 78.922% | 3.980% | 11.411% |
| Mouse Testis PIWIL1-RIP-2 | 0.249% | 0.377% | 2.334% | 2.574% | 0.163% | 78.817% | 4.007% | 11.479% |
| WT_total_smRNA | 73.888% | 5.922% | 4.299% | 0.658% | 5.722% | 1.143% | 3.210% | 5.212% |
| KO-control_total_smRNA | 74.633% | 5.644% | 4.171% | 0.645% | 5.628% | 1.043% | 3.115% | 5.120% |
| WT_PIWIL1-RIP | 1.093% | 1.806% | 63.126% | 5.609% | 0.889% | 0.292% | 5.205% | 21.979% |
| KO-control_PIWIL1-RIP | 1.103% | 1.811% | 63.005% | 5.634% | 0.894% | 0.260% | 5.229% | 22.065% |
| WT_NaIO4-Oxidation | 6.900% | 12.762% | 10.737% | 5.052% | 51.523% | 1.041% | 4.061% | 7.981% |
| KO-#20_total_smRNA | 74.089% | 4.510% | 5.027% | 0.711% | 4.788% | 0.387% | 6.207% | 4.280% |
| KO-#37_total_smRNA | 74.320% | 4.402% | 4.967% | 0.712% | 4.814% | 0.702% | 6.149% | 3.934% |
| KO-#20_PIWIL1-RIP | 0.497% | 1.024% | 70.115% | 4.224% | 1.622% | 0.286% | 4.375% | 17.857% |
| KO-#37_PIWIL1-RIP | 0.481% | 1.026% | 70.168% | 4.225% | 1.634% | 0.369% | 4.364% | 17.733% |
| KO-#20_NaIO4-Oxidation | 10.909% | 19.564% | 18.346% | 3.226% | 30.123% | 1.258% | 6.558% | 10.017% |

**Table S4** Primers used in quantitative RT-PCR.

| Gene | Forward Primer | Reverse Primer |
|---|---|---|
| ACTB | CCCTGGAGAAGAGCTACGAG | GGAAGGAAGGCTGGAAGAGT |
| PIWIL1 | CAGCCAAGTCACAAGGACTC | GATGTCAGCCGGAAATGGTT |
| PIWIL2 | TTGGTTGGAGTAGGACGCTT | AAGGTACAGGGAGGCTTGTC |
| PIWIL4 | GATGGCACCGAGATCACCTA | TGGTCAGTCAGCCCTGTTAG |
| UPF1 | ATCCGCCTGCAGGTCCAGTA | GATCCGCTGCAGTGACACCA |
| FLNA | GGGATCCCATCCCTAAGAGC | CTTGGATGCCACTTTGCCTT |
| LAMC3 | ACTGTGAGCACTGTCAGGAA | TGCAAGGCATGCATTTGTCT |
| CCNE2 | GGGGGATCAGTCCTTGCATT | AAGGCAGCAGCAGTCAGTAT |
| ORC6 | ATGCTGAGGAAAGCAGAGGA | TTCATCCAGGAAGCTGCAAG |
| CDC25A | GTGCCGGTATGTGAGAGAGA | TGCGGAACTTCTTCAGGTCT |
| MCM2 | CGAGATAGAGCTGACTGGCA | CAGTGGCAAAGACAGGGAAG |
| CCND3 | ATCACTGGCACTGAAGTGGA | TCTGTAGGAGTGCTGGTCTG |
| PCNA | GCGTGAACCTCACCAGTATG | TCTCCTGGTTTGGTGCTTCA |
| MYO18B | GAAGCAGATGCACCAGAAGG | CCAATCTGGTCACAGAGGGT |
| VCL | GACCGGCCAAAGCAGCTGTA | GATGGCAGCCTGACCGACTC |
| SESN2 | GGCTGGAGGCACTGATGTCC | TAGTCAGGGTGCAGGCCCAT |
| SRCIN1 | ACAGAGCTCAAGGCTCACTT | GGCTCCTCCTTCAGGAACTT |
| TPM2 | GATGCTGAAGCTGGACAAGG | GGTCCTCAGCTTGCTTCTTG |
| DDIT4 | GAGTCCCTGGACAGCAGCAA | CAGCAGCTGCATCAGGTTGG |
| CCDC80 | AGCAGAAGAAGGAGGGCATT | AGATCACCAGCAACCTCCTC |
| PTEN | GCGGAACTTGCAATCCTCAG | GAACTTGTCTTCCCGTCGTG |
| GADD45G | AGCTGCTGGTTGATCGCACT | AGCAACTCATGCAGCGCTTT |
| GADD45B | TGAATGTGGACCCAGACAGC | AAGGACTGGATGAGCGTGAA |
| TSC2 | TACTGCGTCTGCGACTACAT | CCAGTCAGACTCCTGCTTCA |
| TP53 | GTCCAGATGAAGCTCCCAGA | CAAGAAGCCCAGACGGAAAC |
| MCM10 | GTGGAAGCCTTCTCTGGTCT | CAGCTTCTCTCTGGCCATCT |

**Table S5** Primers used for genome PCR analysis of PIWIL1-knockout SNU-1 cells and cloning of PIWIL1 mutant cDNA.

| Primers ID | Sequence(5'to3') |
|---|---|
| 2U6 hPiwil-1 E5 sg2 For | atgCGTCTCaACCGcattgactataacccactgagttttagagctagaaatagcaag |
| 2U6 hPiwil-1 E5 sg1 Rev | atgCGTCTCgAAACgctgacatcccgtccccagtCGGTGTTTCGTCCTTTCCACAAG |
| hPiwil-1 E5 C9 (genomePCR) For | ACTGTGCCTGCCTTGTCAT |
| hPiwil-1 E5 C9 (genomePCR) Rev | GGTAGTTCTGCTTTGAAGTGGAA |
| PIWIL1-piRNA-binding mutant For | GCAAAATACCTGTGTACAGATTGCCCTACCCCAAGTGCATGTGTGGTGGCCCGAACCTTAGGCAAACAGCAAACTGTCATGGCCATTGCTACAAAGATTGCCCTAGCAATGAACT |
| PIWIL1-piRNA-binding mutant Rev | AATAGCATCGTATTTGTCCTTCCGATTACT |
| (2-250)/(2-624)-F' | gatgatgacgacaaaggatccACTGGGAGAGCCCGAGCC |
| (2-250)-R' | tcaggAGGCCAAATCACCAACCTGTG |
| (384-861)/(384-624)-F' | ttggtgatttggcctCCTGAGCTCTGCTATCTTACAGGTC |
| (384-861)/(759-861)-R' | aacgggccctctagactcgagTTAGAGGTAGTAAAGGCGGTTTGA |
| (2-624)/(384-624)-R' | tgttccaggaagtggCTTCAGGGGGATGTCCACCC |
| (759-861)-F' | tgaagCCACTTCCTGGAACAGTTATTGATG |
